# RTT109 and Fun30 proteins mediate epigenetic regulation of the DNA damage response pathway in *C. albicans*

**DOI:** 10.1101/2021.07.22.453354

**Authors:** Pramita Garai, Prashant Kumar Maurya, Himanshu Bhatt, Aarti Goyal, Sakshi Dewasthale, Meghna Gupta, Dominic Thangminlen Haokip, Rohini Muthuswami

**Affiliations:** Chromatin Remodelling Laboratory, School of Life Sciences, JNU, New Delhi 110067

**Keywords:** Rtt109, Fun30, DNA damage response, Epigenetics

## Abstract

Fun30, an ATP-dependent chromatin remodeller, from *S. cerevisiae* mediates both regulation of gene expression as well as DNA damage response/repair. In this paper, we have characterized the biochemical and physiological function of Fun30 from the opportunistic fungi, *C. albicans*. Biochemically, the protein shows DNA-stimulated ATPase activity. Physiologically, the protein co-regulates transcription of *RTT109*, *TEL1*, *MEC1*, and *SNF2*-genes that encode for proteins involved in DNA damage response and repair pathway. The expression of *FUN30*, in turn, is regulated by histone H3 acetylation catalysed by Rtt109 encoded by *RTT109*. The *RTT109Hz/FUN30Hz* mutant strain shows sensitivity to oxidative stress and resistance to MMS as compared to the wild type strain. Quantitative PCR showed that the sensitivity to oxidative stress results from downregulation of *MEC1*, *RAD9*, *MRC1* and *RAD5* expression; ChIP experiments showed Fun30 but not H3ac regulates the expression of these genes in response to oxidative stress. In contrast, on treatment with MMS, the expression of *RAD9* is upregulated and this upregulation is co-regulated by both Fun30 and H3 acetylation catalysed by Rtt109. Thus, Fun30 and H3 acetylation mediate the response of the fungal cell to genotoxic agents in *C. albicans* by regulating the expression of DNA damage response and repair pathway genes.

## INTRODUCTION

The ATP-dependent chromatin remodelling proteins modulate gene expression as well as mediate DNA damage response/repair by using the energy released from ATP hydrolysis (1). Though classified as helicases and possessing the seven conserved helicase motifs, these proteins do not possess the canonical helicase activity (2, 3). Instead, these proteins use the energy stored in ATP to reposition, evict, or slide nucleosomes (4). In addition, they can also mediate histone variant exchange (5).

The ATP-dependent chromatin remodelling proteins are classified into 24 sub-families of which the Etl1 sub-family comprises of Etl1 in mouse, SMARCAD1 in human, Fun30 in *S. cerevisiae*, and Fft1, Fft2, Fft3 in *S. pombe* (2). All the members of the sub-family have been shown to mediate DNA end resection during double-strand break repair, thus, indicating that these proteins are involved in the repair process (6, 7). Studies have also shown that these proteins are required for maintaining the chromatin structure at heterochromatic loci (8, 9). Biochemical studies have shown that the Fun30 from *S. cerevisiae* is a homodimer possessing DNA-stimulated ATPase activity (10). These studies have also shown that the purified protein can mediate nucleosome sliding as well as histone variant exchange; however, the efficiency in mediating histone variant exchange is greater than nucleosome sliding (10). The protein has not yet been characterized in *C. albicans*.

In addition to ATP-dependent chromatin remodelling proteins, histone modifying enzymes have also been shown to play a role in DNA repair. Rtt109 is a B-type histone acetyltransferase specific to yeast and fungi (11). Studies have shown that the protein acetylates histone H3 at K56, K9, and K27 (12). The H3K56ac modification is associated with DNA damage response both in *S. cerevisiae* and in *C. albicans* (13, 14). This modification has been recently identified in mammalian cells too where p300 has been shown to mediate the acetylation at K56 position of histone H3 (15). Studies have shown that Rtt109 and p300 do not share sequence homology but do share structural similarity (16).

As both Rtt109 and Fun30 are involved in DNA repair, therefore, in this paper we seek to ask whether there is any crosstalk between Fun30 and Rtt109 in *C. albicans*. *C. albicans* is an opportunistic pathogen causing candidiasis in immuno-compromised patients. The organism is an intracellular pathogen and the host cell combats the infection by generating ROS that causes DNA damage. Thus, *C. albicans* has evolved robust mechanisms to repair the damaged DNA thereby ensuring genomic stability (17). As stated earlier, the role of Rtt109 but not of Fun30 has been delineated in *C. albicans* (14). Rtt109 in *C. albicans* has been linked to pathogenesis; the cells lacking Rtt109 were found to be sensitive to genotoxic agents like camptothecin (CPT) and methyl methane sulfonate (MMS) indicating the role of the histone acetyltransferase in DNA damage response/repair (14). Recently, we have shown that Rtt109 mediated acetylation of H3 regulates the expression of genes involved in GPI anchor biosynthesis and in ergosterol biosynthesis (18). However, the crosstalk between Rtt109 and Fun30 has not yet been investigated.

In this paper we have characterized the biochemical and functional activities of Fun30 from *C. albicans*. We show that Fun30 regulates the expression of Rtt109 and in turn, Rtt109 regulates the expression of Fun30 via H3 acetylation (H3ac). Fun30 and H3ac are needed for the expression *TEL1* and *MEC1* encoding for the sensor kinases, Tel1 and Mec1 respectively. Finally, we show that Fun30 regulates the expression of Rad9, a cell cycle checkpoint protein, and Rad5, an ATP-dependent chromatin remodelling protein, in response to genotoxic stress, thus, modulating the DNA damage response pathway on induction of DNA damage.

## RESULTS

### The orf19.6291 encodes for Fun30, a member of the ATP-dependent chromatin remodelling protein family

*C. albicans* orf19.6291 is an uncharacterized protein annotated as Fun30 in the Candida Genome database (19). Phylogenetic analysis showed that orf19.6291 is closely related to Fun30 from *S. cerevisiae* (Supplementary Fig. 1A). Further, bootstrap values suggest that the orf19.6291 evolved along with Fft3, Fft2, SMARCAD1 and Etl1 (Supplementary Fig. 1A). Sequence analysis showed that orf19.6291 contained the conserved helicase motifs as well as CUE motif (Supplementary Figs. 1B and 2). The CUE motif that has been shown to recognize and bind to monoubiquitinated proteins (20) is present in the N-terminus of Fun30 in case of *S. cerevisiae*. However, the CUE motif in orf19.6291, as in the case of Fft2, is present between the subdomains of the helicase motifs (Supplementary Fig. 1B). Based on these analyses, it was concluded that orf19.6291 encodes for Fun30 in *C. albicans*.

### Fun30 localizes to the nucleus

To understand the localization of Fun30 in *C. albicans*, one of the alleles of *FUN30* in SN152 strain was myc-tagged at the C-terminus as explained in the Materials and Methods section. This strain is, hereinafter, referred as *FUN30myc*. Using anti-myc antibody, the localization of the protein was found to be predominantly in the nucleus (Supplementary Fig. 1C).

### Fun30 in *C. albicans* is a DNA-stimulated ATPase with fork DNA being the optimal effector

To biochemically characterize Fun30 from *C. albicans* we attempted to purify the full-length protein by overexpressing it in *E. coli*. However, the yield of the protein was not sufficient to perform biochemical assays. Therefore, a truncated version of the protein lacking the N-terminus from amino acids 1 to 512 (ΔNFun30) was cloned and overexpressed in *E. coli* (Supplementary Fig. 1D and Fig. 1A). The purified protein showed DNA-independent activity like *S. cerevisiae* Fun30 (Fig. 1B). The ATPase activity was stimulated ~1.5 fold in the presence of the 500 nM fork DNA while other DNA molecules did not yield any significant stimulation (Fig. 1B). The K_M_ for the interaction with fork DNA was calculated to be 0.06 ± 0.007 μM (Fig. 1C). The binding parameters, using fluorescence spectroscopy, for the interaction of ATP and DNA with the protein were calculated both in the absence and presence of the other ligand. The binding studies showed that the interaction of fork DNA with the protein was similar both in the absence and presence of saturating concentration (60 μM) of ATP (Fig. 1D and F). In contrast, the interaction of ATP was 2-fold weaker in the presence of saturating concentration (6 μM) of fork DNA (Fig. 1E and F).

**Figure 1.**
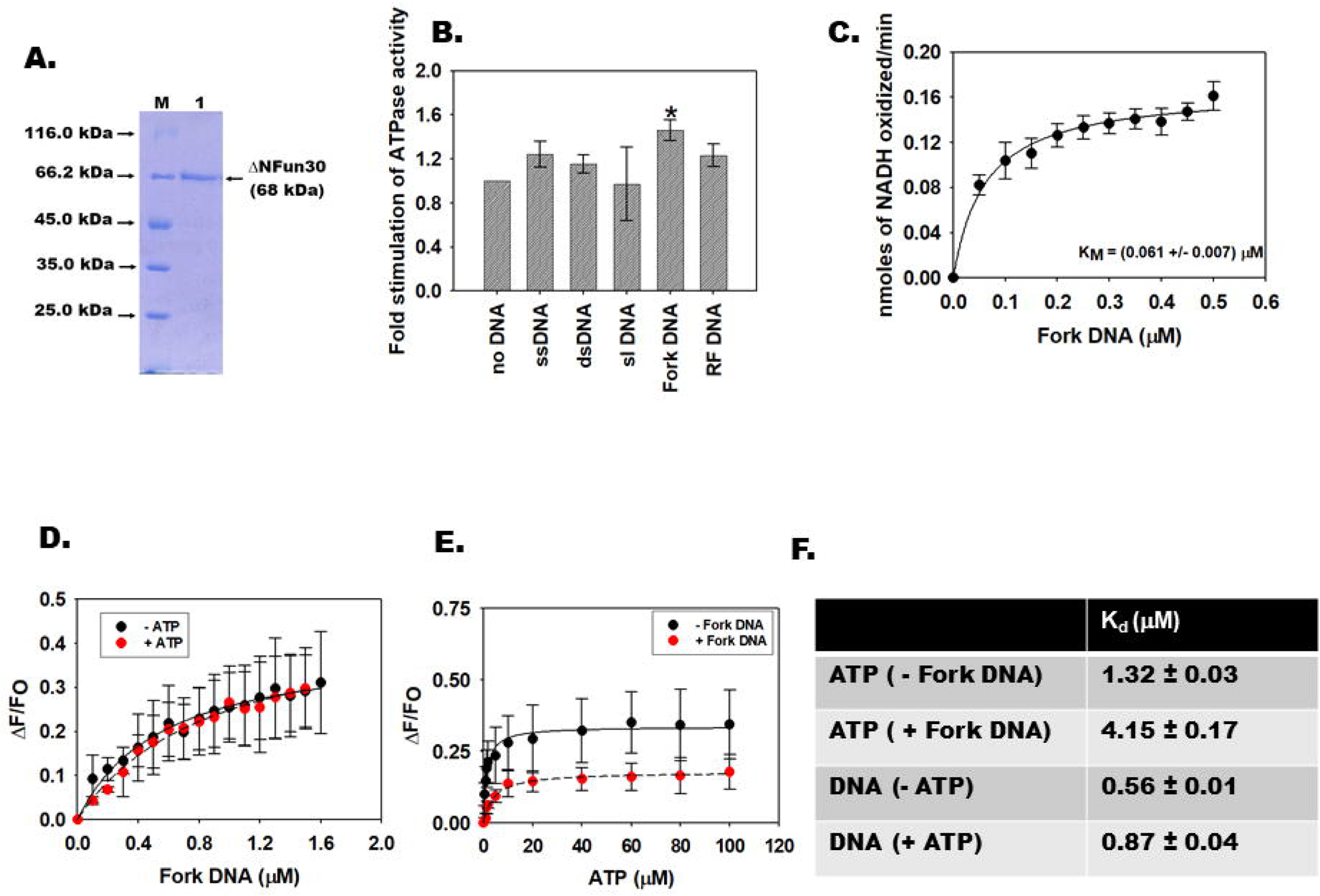
Fun30 in *C. albicans* is a DNA-stimulated ATPase with fork DNA being the optimal effector. (A). Purification of recombinant ΔNFun30 from *E. coli*. (B). ATPase activity was performed as explained in Materials and Methods using different DNA effector molecules. The protein concentration was 200 nM and the DNA concentration was 500 nM in these experiments. The protein shows DNA-independent ATPase activity that is stimulated in the presence of Fork DNA. (C). The K_M_ was calculated for Fork DNA by estimating ATPase activity in the presence of increasing concentration of DNA. The protein concentration was 200 nM in this experiment. (D). Binding constant (K_d_) was calculated using fluorescence spectroscopy for the interaction of ΔNFun30 with DNA in the absence and presence of 60 μM ATP. (E). Binding constant (K_d_) was calculated using fluorescence spectroscopy for the interaction of ΔNFun30 with ATP in the absence and presence of 6 μM Fork DNA. (F). Table depicting the K_d_ values for the interaction of ATP and Fork DNA with ΔNFun30.The K_d_ and K_M_ are expressed as average ± s.d. of three independent experiments.

### Fun30 regulates the expression of *RTT109*, *SNF2, TEL1*, and *MEC1* in *C. albicans*

To understand the physiological role of Fun30 in *C. albicans* a heterozygote mutant was created where one copy of the gene was deleted in SN152 as well as in BWP17 strain of *C. albicans*. In case of SN152, we used the *FUN30myc* strain where one copy of *FUN30* was fused with *myc* to encode Fun30myc protein.

The heterozygous mutant was confirmed using PCR and *HIS1* cassette specific primers in both *FUN30myc* and BWP17 strains (Supplementary Fig. 3A-C). Hereinafter, the heterozygous mutant, *FUN30/fun30,* is referred as *FUN30mycHz* if made in *FUN30myc* background and as *FUN30Hz* if made in BWP17 background. We ensured that in *FUN30mycHz* strain, the copy lacking the *myc* tag was deleted to create the *FUN30mycHz* mutant. Quantitative PCR (qPCR) showed that the expression of *FUN30* was downregulated in the heterozygote mutant both in *FUN30myc* and BWP17 strains (Fig. 2A and B). We were unable to create homozygous null mutant, *fun30/fun30*. We tried creating homozygous null mutant using *ARG4* marker both in *FUN30myc* and BWP17 strains. We made at least 6 attempts for each strain screening more than 120 colonies, but we did not get any null mutant. Supplementary Fig. 3D shows the gel picture of one such attempt made in *FUN30Hz* strain. We also tried to make a conditional null mutant using *MET3* promoter in BWP17 strain. We made 4 attempts and screened more than 20 colonies. Once again, we were unsuccessful. However, both copies could be deleted when *FUN30* was ectopically expressed in BWP17 (Supplementary Fig. 3E and F). For this experiment, *FUN30* was cloned into p*ACT1* vector and overexpressed in BWP17 using *URA3* as selection marker. One copy of *FUN30* was deleted using *HIS1* specific cassette and the other copy was deleted using *ARG4* specific cassette. As *FUN30myc* possesses *URA3*, this experiment could not be performed in this strain.

**Figure 2.**
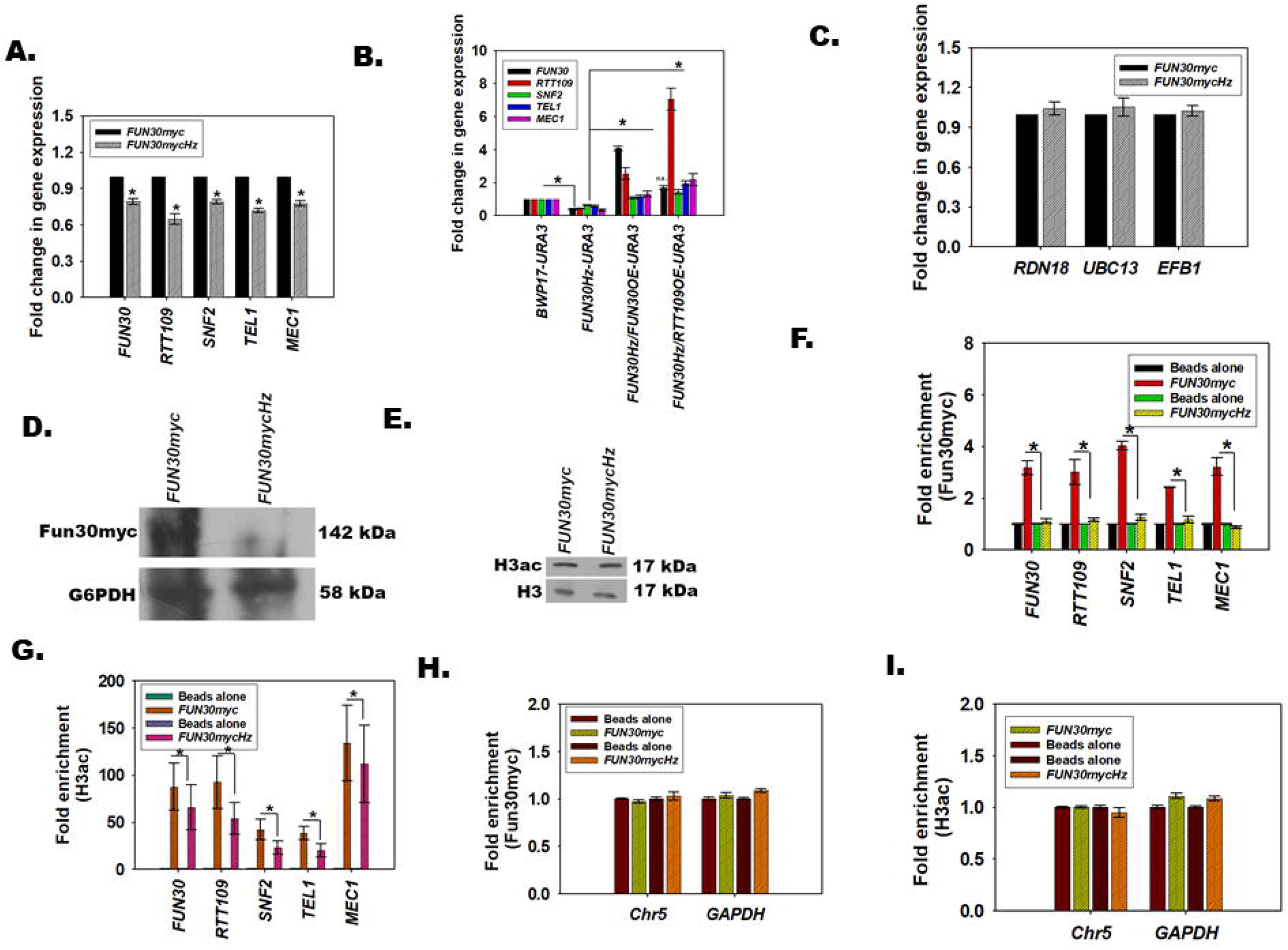
Fun30 regulates the expression of *RTT109*, *SNF2, TEL1,* and *MEC1* in *C. albicans*. (A). Expression of *FUN30, RTT109, SNF2, TEL1,* and *MEC1* in *FUN30mycHz* was compared to the expression in *FUN30myc* strain using qPCR. (B). Expression of *FUN30, RTT109, SNF2, TEL1,* and *MEC1* in analysed in BWP17-*URA3*, *FUN30Hz*-*URA3, FUN30Hz/FUN30OE-URA3,* and, *FUN30Hz/RTT109OE-URA3* using qPCR. (C). Expression of *RDN18*, *UBC13* and *EFB1* was analysed by qPCR in *FUN30myc* and *FUN30mycHz* strain. (D). Expression of Fun30myc was analysed by western blot in *FUN30myc* and *FUN30mycHz* strains. G6PDH was used as internal control. (E). Expression of H3ac was analysed by western blot in *FUN30myc* and *FUN30mycHz* strains. H3 was used as internal control. (F). The occupancy of Fun30myc on *FUN30, RTT109, SNF2, TEL1,* and *MEC1* promoters was compared in *FUN30myc* and *FUN30mycHz* strains by ChIP. (G). The occupancy of H3ac on *FUN30, RTT109, SNF2, TEL1,* and *MEC1* promoters was compared in *FUN30myc* and *FUN30mycHz* strains by ChIP. (H). The occupancy of Fun30myc on the intergenic regions of Chr5 and on *GAPDH* promoter was compared in *FUN30myc* and *FUN30mycHz* strains by ChIP. (I). The occupancy of H3ac on the intergenic regions of Chr5 and on *GAPDH* promoter was compared in *FUN30myc* and *FUN30mycHz* strains by ChIP. The qPCR and ChIP data are presented as average ± s.e.m. of three biological replicates. Star indicates p <0.05. The western blots were repeated for three biological replicates and one representative image is shown.

Previously, we had shown that in mammalian cells the DNA damage response pathway is transcriptionally regulated by ATP-dependent chromatin remodelling proteins (21–23). Fun30, an ATP-dependent chromatin remodelling protein, mediates not only DNA end resection but also modulates transcriptional co-regulation (7, 24–26). Therefore, we investigated whether Fun30 regulates genes involved in the DNA damage response pathway in *C. albicans*. Specifically, we sought to investigate the expression of Rtt109 that acetylates H3 at K56 and other positions, Tel1 and Mec1 that are functional homologs of ATM and ATR, and Snf2 that is the functional homolog of BRG1. Tel1 and Mec1, like ATM and ATR, are kinases that are activated on DNA damage and transduce the signal to downstream effectors via phosphorylation (27). In mammalian cells, BRG1 has been shown to not only modulate the expression of ATM and ATR but also to directly participate in DNA damage repair (21, 28, 29).

Quantitative PCR analysis showed that the expression of *RTT109*, *SNF2*, *TEL1*, and *MEC1* was downregulated in *FUN3mycHz* as well as *FUN30Hz* strains (Fig. 2A and B). The expression of these genes was restored when *FUN30* was overexpressed in *FUN30Hz* background ectopically under the regulation of *ACT1* promoter (Fig. 2B). Interestingly, the expression of the genes was also restored when *RTT109* was overexpressed under the regulation of constitutive *ACT1* promoter in *FUN30Hz* background (Fig. 2B).

To show Fun30 specifically regulates the expression of a subset of genes, the expression of three genes-*RDN18* (encodes for 18S ribosomal RNA), *EFB1* (encodes translation elongation factor beta), and *UBC13* (encodes for ubiquitin conjugating enzyme) - was analysed. The expression of these genes was unchanged in *FUN30mycHz* as compared to the *FUN30myc* wild type strain (Fig. 2C).

Western blots using anti-myc antibody confirmed that the expression of Fun30myc was reduced in the whole cell lysate of *FUN30mycHz* strain (Fig. 2D). However, the levels of H3ac (probed using anti-H3K56ac antibody) was not altered in the whole cell lysate of *FUN30mycHz* (Fig. 2E). It needs to be pointed out that the antibody against H3K56ac is non-specific and can also recognize H3K9ac (30), and therefore, we can only conclude that though the transcript levels of *RTT109* are downregulated, the total H3ac in the whole cell lysate is not altered.

### Fun30 binds to the promoter regions of the DNA damage response genes

To assess whether Fun30 directly regulates the expression of *RTT109*, *SNF2*, *TEL1* and *MEC1* genes, ChIP assay was performed using anti-myc antibody to probe the interaction of Fun30myc protein with the promoter regions of these genes. In addition, the interaction of Fun30myc protein with the *FUN30* promoter was also investigated to understand whether Fun30 regulates its own expression. Experimental results confirmed that Fun30 did bind to the promoter regions of these genes (Fig. 2F). In addition, the protein also bound to its own promoter indicating that it may be regulating itself (Fig. 2F). The occupancy of H3ac on the promoter regions of *FUN30*, *RTT109*, *SNF2*, *TEL1*, and *MEC1* was found to be also reduced indicating that the global levels of H3ac does not alter yet its recruitment to the promoter is affected in *FUN30mycHz* mutant strain (Fig. 2G). The occupancy of Fun30 and H3ac on intergenic regions present in Chr5 as well as on *GAPDH* promoter was not altered indicating Fun30 specifically localizes to the promoter regions of the DNA damage response genes and regulates their expression (Fig. 2H and I).

### H3ac regulates the expression of *FUN30, SNF2*, *TEL1*, and *MEC1*

As the above experiments showed that H3ac is present on the promoter of DNA damage response genes and that overexpression of *RTT109* restored the expression of the genes including *FUN30*, we asked whether a feedback loop exists in *C. albicans* wherein H3ac catalysed by Rtt109 regulates *FUN30* in a manner similar to the feedback loop between SMARCAL1 and BRG1 in mammalian cells (23). The expression of *FUN30* along with *SNF2*, *TEL1* and *MEC1* was analysed in cells where both copies of *RTT109* was deleted to create homozygous null mutant, *rtt109/rtt109* (hereafter termed as *RTT109D*), using PCR based strategies (Supplementary Fig. 4A and B). A revertant strain, *rtt109/rtt109/*p*ACT1-RTT109* (*RTT109D*/*RTT109OE-URA3*), was also created wherein *RTT109* was cloned into p*ACT1* vector and integrated into *RPS1* locus. As this strain in BWP17 was created using *URA3* as the marker, all comparisons were done using BWP17-*URA3* strain created specifically for this purpose. In *FUN30myc*, the two copies of *RTT109* was deleted using *HIS1* and *ARG4* markers (Supplementary Fig. 4C).

Quantitative PCR showed that the expression *FUN30*, *SNF2*, *TEL1*, and *MEC1* was downregulated in *RTT109D* as compared to the wild type irrespective of the strain background (Fig. 3A and B). This downregulation was specific as *RDN18*, *UBC13*, and *EFB1* expression was unaltered (Supplementary Fig. 4D). Overexpression of *RTT109* under the control of *ACT1* promoter in *RTT109D* background (*RTT109D*/*RTT109OE-URA3*) restored the expression level of these genes (Fig. 3A). Western blots confirmed that H3ac levels were downregulated in *RTT109D* mutant and restored back in *RTT109D*/*RTT109OE-URA3* strain (Fig. 3C). Interestingly, Fun30 expression was also not observed in *RTT109D* mutant (Fig. 3D).

**Figure 3.**
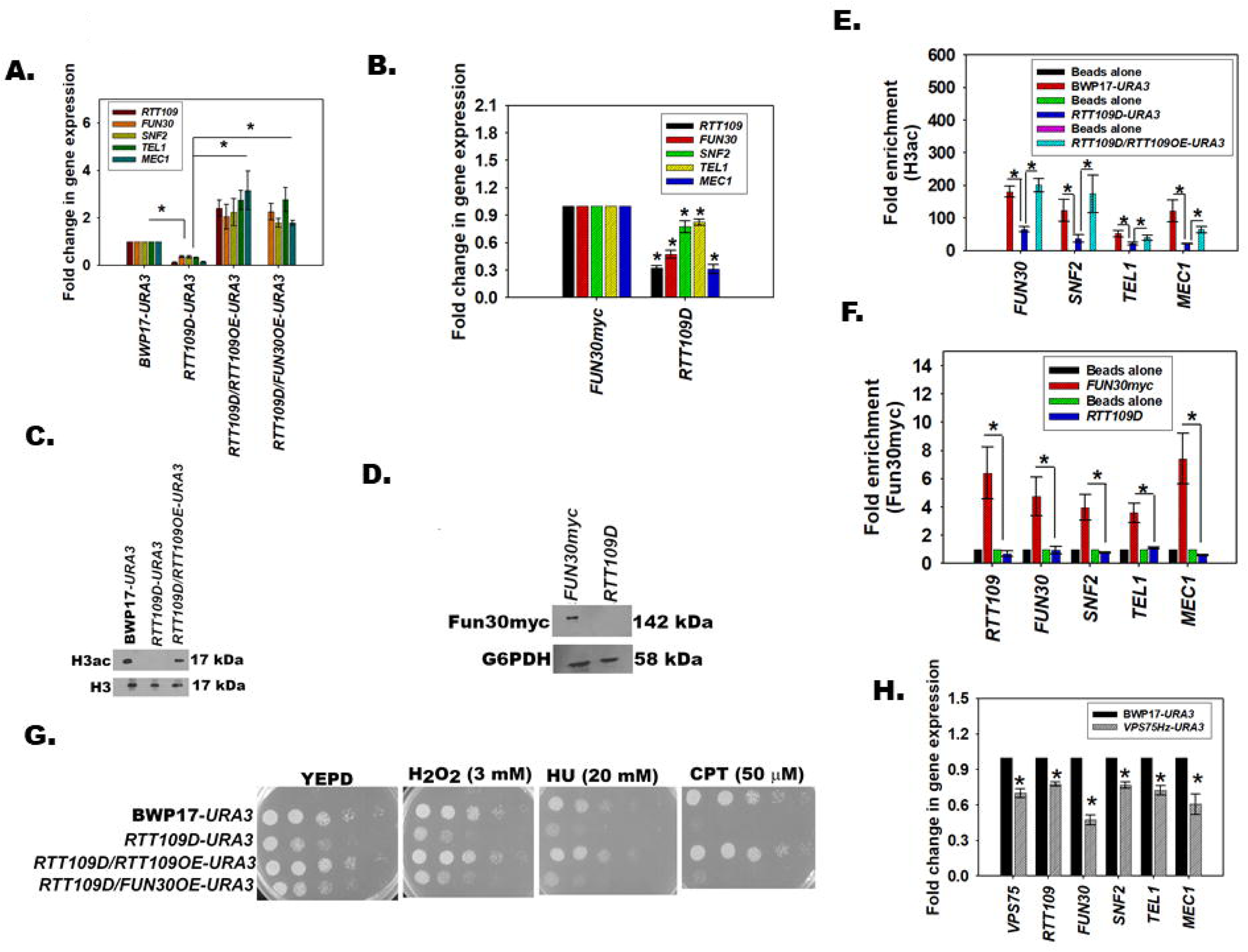
H3ac regulates the expression of *FUN30, SNF2*, *TEL1*, and *MEC1*. (A). Expression of *RTT109, FUN30, SNF2, TEL1,* and *MEC1* were analysed in BWP17-*URA3*, *RTT109D*-*URA3, RTT109D/RTT109OE-URA3,* and *RTT109D/FUN30OE-URA3* using qPCR. (B). Expression of *RTT109, FUN30, SNF2, TEL1,* and *MEC1* were analysed in *FUN30myc* and *RTT109D* strains using qPCR. (C). Expression of H3ac was analysed by western blot in BWP17*-URA3*, *RTT109D*-*URA3, RTT109D/RTT109OE-URA3* strains. H3 was used as internal control. (D). Expression of Fun30myc was analysed by western blot *FUN30myc* and *RTT109D* strains. G6PDH was used as internal control. (E). Occupancy of H3ac on *FUN30, SNF2, TEL1,* and *MEC1* promoters was analysed using ChIP in BWP17-*URA*3, *RTT109D*-*URA3, RTT109D/RTT109OE-URA3* strains. (F). Occupancy of Fun30myc on *FUN30, SNF2, TEL1,* and *MEC1* promoters was analysed using ChIP in *FUN30myc* and *RTT109D* strains. (G). Sensitivity of BWP17-*URA3*, *RTT109D*-*URA3, RTT109D/RTT109OE-URA3,* and *RTT109D/FUN30OE-URA3* to genotoxic stress was observed using plate assays. (H). Expression of *VPS75*, *RTT109*, *FUN30*, *SNF2*, *TEL1*, and *MEC1* was analysed in BWP17-*URA3* and *VPS75Hz-URA3* strains by qPCR. The qPCR and ChIP data are presented as average ± s.e.m. of three biological replicates. Star indicates p <0.05. The western blots were repeated for three biological replicates and one representative image is shown.

ChIP analysis showed that H3ac was present on the promoters of *FUN30*, *SNF2*, *TEL1* and *MEC1* (Fig. 3E). The occupancy of H3ac decreased in *RTT109D* mutant but was restored in *RTT109OE* strain (Fig. 3E). Finally, occupancy of Fun30myc tagged protein was also found to be decreased on the promoters of the DNA damage response genes in *RTT109D* mutant (Fig. 3F).

Thus, from these experimental results, it was concluded that both H3ac, catalysed by Rtt109, and Fun30 regulate the expression of each other as well as of *SNF2*, *TEL1*, and *MEC1*.

### Overexpression of *FUN30* in *RTT109D* mutant restores the gene expression

As the above studies indicated that the expression of *FUN30* and *RTT109* was co-regulated, therefore, we next asked whether overexpression of *FUN30* in *RTT109D* can restore the gene expression and therefore, the defects observed in mutant strain. Therefore, *FUN30* was overexpressed under the *ACT1* promoter in *RTT109D* background (*RTT109D*/*FUN30OE-URA3).* Quantitative PCR showed that the expression of *SNF2, TEL1,* and *MEC1* was restored in this strain; however, the sensitivity to DNA damage agents was not restored (Fig. 3A and G). It has already been shown in *S. cerevisiae* that H3K56ac deposition on the newly replicated DNA strand is important for completion of DNA repair (31). Therefore, even though overexpression of Fun30 restores gene expression, DNA repair most probably continues to be impaired due to the absence of H3K56ac.

### The catalytic activity of Rtt109 is essential for the transcription regulation

The catalytic activity of Rtt109 requires the help of two chaperones-Asf1 and Vps75 (32). Therefore, we hypothesized that blocking the catalytic activity of Rtt109 by deleting *VPS75* would also result in downregulation of *RTT109*, *FUN30*, *SNF2*, *TEL1*, and *MEC1* expression. To test the hypothesis, one copy of Vps75 was deleted (*VPS75/vps75* referred hereinafter as *VPS75Hz*) using PCR based strategy in BWP17 strain. Quantitative PCR confirmed that the expression of *RTT109*, *FUN30*, *SNF2*, *TEL1*, and *MEC1* was indeed downregulated (Fig. 3H) in the mutant strain confirming that the catalytic activity of Rtt109 was essential for the transcriptional regulation mediated by H3ac.

### *RTT109Hz/FUN30Hz* double mutant shows differential response to genotoxic stress

To understand how Rtt109 and Fun30 regulate the DNA damage response pathway in the presence of genotoxic stress, double heterozygous mutant wherein one copy each of *RTT109* and *FUN30* (*RTT109/rtt109;FUN30/fun30*, hereinafter termed as *RTT109Hz*/*FUN30Hz*) was deleted BWP17. A similar mutant was made in *FUN30myc* strain and termed as *RTT109Hz/FUN30mycHz*. Quantitative PCR confirmed that the expression of *RTT109*, *FUN30, SNF2, TEL1*, and *MEC1* was downregulated in both *RTT109Hz/FUN30mycHz* and *RTT109/FUN30Hz* strains (Fig. 4A and B). In contrast, the expression of *RDN18*, *UBC13*, and *EFB1* was unaltered (Fig. 4C). The ability of the *RTT109Hz*, *FUN30Hz*, *RTT109Hz/FUN30mycHz* and *RTT109Hz/FUN30Hz* to grow in the presence of H_2_O_2_ (generates ROS and induces base modifications), CPT (results in double-strand breaks) and MMS (induces methylation of bases) was studied using plate (spot) assays. As compared to the wild type strain, both *RTT109Hz/FUN30mycHz* and *RTT109Hz/FUN30Hz* mutant showed no response to CPT (Fig. 4D and E), were sensitive to H_2_O_2_ (Fig. 4D and E) and resistant to MMS (Fig. 4D and E).

**Figure 4.**
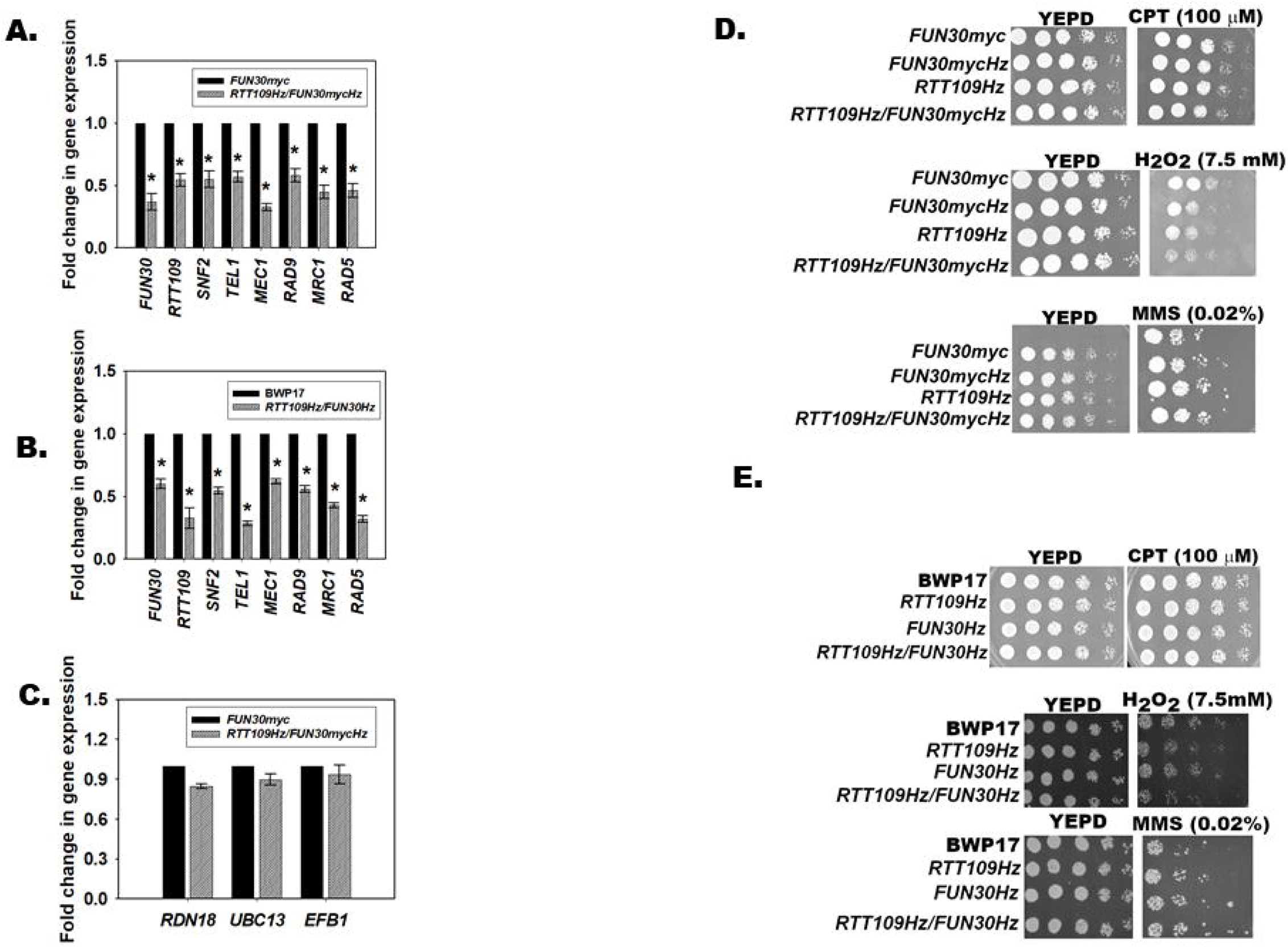
*RTT109Hz/FUN30Hz* double mutant shows differential response to genotoxic stress. (A). Expression of *FUN30*, *RTT109*, *SNF2*, *TEL1*, *MEC1*, *RAD9*, *MRC1*, and *RAD5* was evaluated in *FUN30myc* and *RTT109Hz/FUN30mycHz* strains using qPCR. (B). Expression of *FUN30*, *RTT109*, *SNF2*, *TEL1*, *MEC1*, *RAD9*, *MRC1*, and *RAD5* was evaluated in BWP17 and *RTT109Hz/FUN30Hz* strains using qPCR. (C). Expression of *RDN18*, *UBC13*, and *EFB1* was analysed by qPCR in *RTT109Hz*/*FUN30mycHz* strain. (D). Sensitivity of *FUN30myc*, *RTT109Hz*, *FUN30Hz*, and *RTT109Hz/FUN30mycHz* strains was studied in the presence of CPT (100 μM), H_2_O_2_ (7.5 mM), and MMS (0.02%). (E). Sensitivity of BWP17, *RTT109Hz*, *FUN30Hz*, and *RTT109Hz/FUN30Hz* strains was studied in the presence of CPT (100 μM), H_2_O_2_ (7.5 mM), and MMS (0.02%). The qPCR data is presented as average ± s.e.m. of three biological replicates. Star indicates p <0.05. Plates were incubated at 30°C and imaged after 24 hrs.

Thus, the response of the mutant strain appears to be dependent on the type of damage induced by the genotoxic agent.

#### Rtt109 and Fun30 co-regulate the transcription of genes involved in DNA damage response pathways

Hydrogen peroxide, CPT and MMS activate base excision repair, double-strand repair, and mismatch repair pathway respectively (33–35). The Mec1 kinase is a central transducer in these pathways wherein after activation by phosphorylation it phosphorylates Rad9 and Mrc1. Rad9 functions in the DNA damage checkpoint (DDC) while Mrc1 is important for the DNA replication checkpoint (DRC) (36). Both proteins promote phosphorylation of Rad53, also known as effector kinase, by Mec1, thus, activating the cell cycle checkpoint. In addition, Rad5, an ATP-dependent chromatin remodelling protein as well as E3 ubiquitin ligase, is required for the bypass of replication forks through MMS-damaged DNA (37). In *S. cerevisiae*, Fun30 and Rad5 cooperate to repair the stalled replication fork. In the absence of Rad5, Fun30 null mutants show resistance to MMS (38).

We hypothesized that the DNA damage response is epigenetically regulated by H3ac (catalysed by Rtt109) and Fun30. Further, based on the plate assays, we postulated that in the presence of H_2_O_2_, *MEC1*, *RAD9*, and *MRC1* would be downregulated in *RTT109Hz/FUN30Hz* strain as compared to the wild type strain; in the presence of CPT, the expression of *MEC1*, *RAD9*, and *MRC1* would be unchanged between the wild type and the mutant strain; and in the presence of MMS, the expression of *RAD5* would be downregulated leading to resistance phenotype.

Therefore, the expression of *MEC1*, *RAD9*, *MRC1*, and *RAD5* along with *FUN30*, *RTT109*, *SNF2*, and *TEL1* was evaluated in the mutant strain (*RTT109/FUN30mycHz* as well as *RTT109/FUN30Hz*) and compared with the wild type strain (*FUN30myc* and BWP17 respectively) in the absence and presence of the genotoxic stress.

Quantitative PCR showed that in the presence of H_2_O_2_, the expression of *FUN30*, *RTT109*, *SNF2*, *TEL1*, *MEC1*, *RAD9*, *MRC1*, and *RAD5* was downregulated in the mutant strain as compared to the wild type strain, thus, resulting in sensitive phenotype (Fig. 5 and Supplementary Fig. 5).

**Figure 5.**
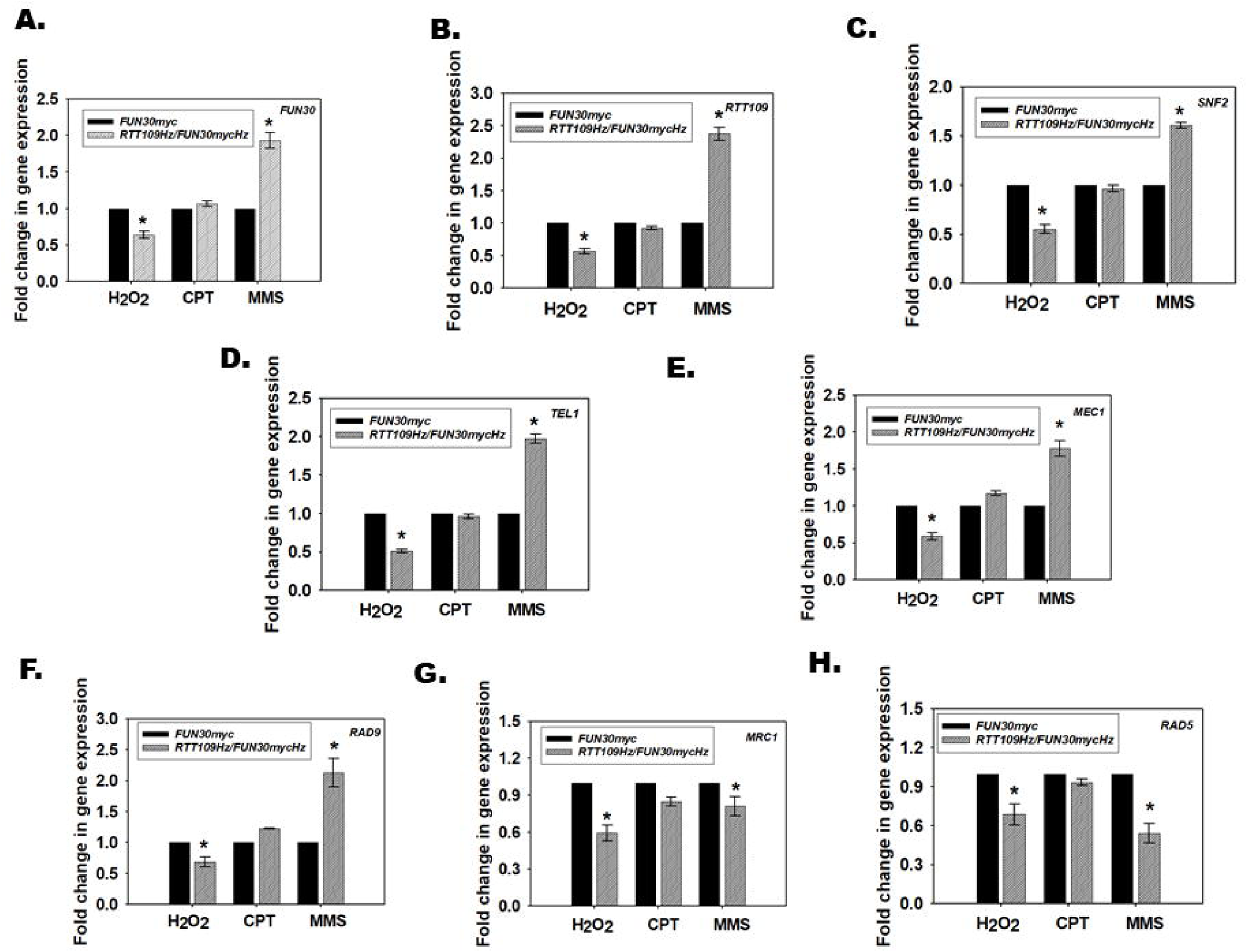
Rtt109 and Fun30 co-regulate the transcription of genes involved in DNA damage response pathways. Expression of (A). *FUN30*; (B). *RTT109*; (C). *SNF2*; (D). *TEL1*; (E). *MEC1*; (F) *RAD9*; (G). *MRC1*; (H). *RAD5* was analysed in *FUN30myc* and *RTT109Hz/FUN30mycHz* strains in the presence of H_2_O_2_ (7.5 mM), CPT (100 μM) and MMS (0.02%). The qPCR data is presented as average ± s.e.m. of three biological replicates. Star indicates p <0.05.

In contrast, in the presence of CPT, *MEC1* and *RAD9* expression was unchanged in the mutant strain as compared to the wild type strain (Fig. 5 and Supplementary Fig. 5). Interestingly, some strain specific differences were observed. The expression of *FUN30*, *RTT109*, *SNF2*, *TEL1*, *RAD5* and *MRC1* was unchanged in *RTT109/FUN30mycHz* (Fig. 5).

However, the expression of *FUN30*, *RTT109*, and *SNF2* was downregulated while *TEL1* was upregulated in *RTT109Hz/FUN30Hz* (Supplementary Fig. 5D). Taken together, in both strains, as *MEC1* expression is unchanged, we postulate that the DNA damage response is operational in the mutant strain in the presence of CPT, and therefore, no phenotypic response was observed to this stress agent.

In the presence of MMS too, expression of *MEC1* was unchanged but the expression of *TEL1*, *MRC1*, and *RAD5* was downregulated but *RAD9* was upregulated in the mutant strain as compared to the wild type strain (Fig. 5 and Supplementary Fig. 5).

Thus, our data shows that Rtt109 and Fun30 are epigenetically regulating the DNA damage response pathway depending on the type of DNA damage induced by the genotoxic stress.

#### The response to DNA damage is regulated by the occupancy of Fun30 on RAD9, MRC1, and RAD5 promoter

To understand how Rtt109 and Fun30 epigenetically regulate the DNA damage response pathway, ChIP experiments were performed to study the occupancy of H3ac and Fun30 on the promoters of *FUN30*, *RTT109*, *SNF2*, *TEL1*, *MEC1*, *RAD9*, *MRC1*, and *RAD5* genes in *FUN30myc* and *RTT109Hz*/*FUN30mycHz* mutant strain. In untreated cells, the occupancy of both Fun30myc and H3ac decreased on these gene promoters in the *RTT109Hz*/*FUN3mycHz* strain as compared to the *FUN30myc* wild type strain (Fig. 6A and B). The decreased occupancy correlates with decreases expression of these genes in the mutant strain as compared to the wild type strain.

**Figure 6.**
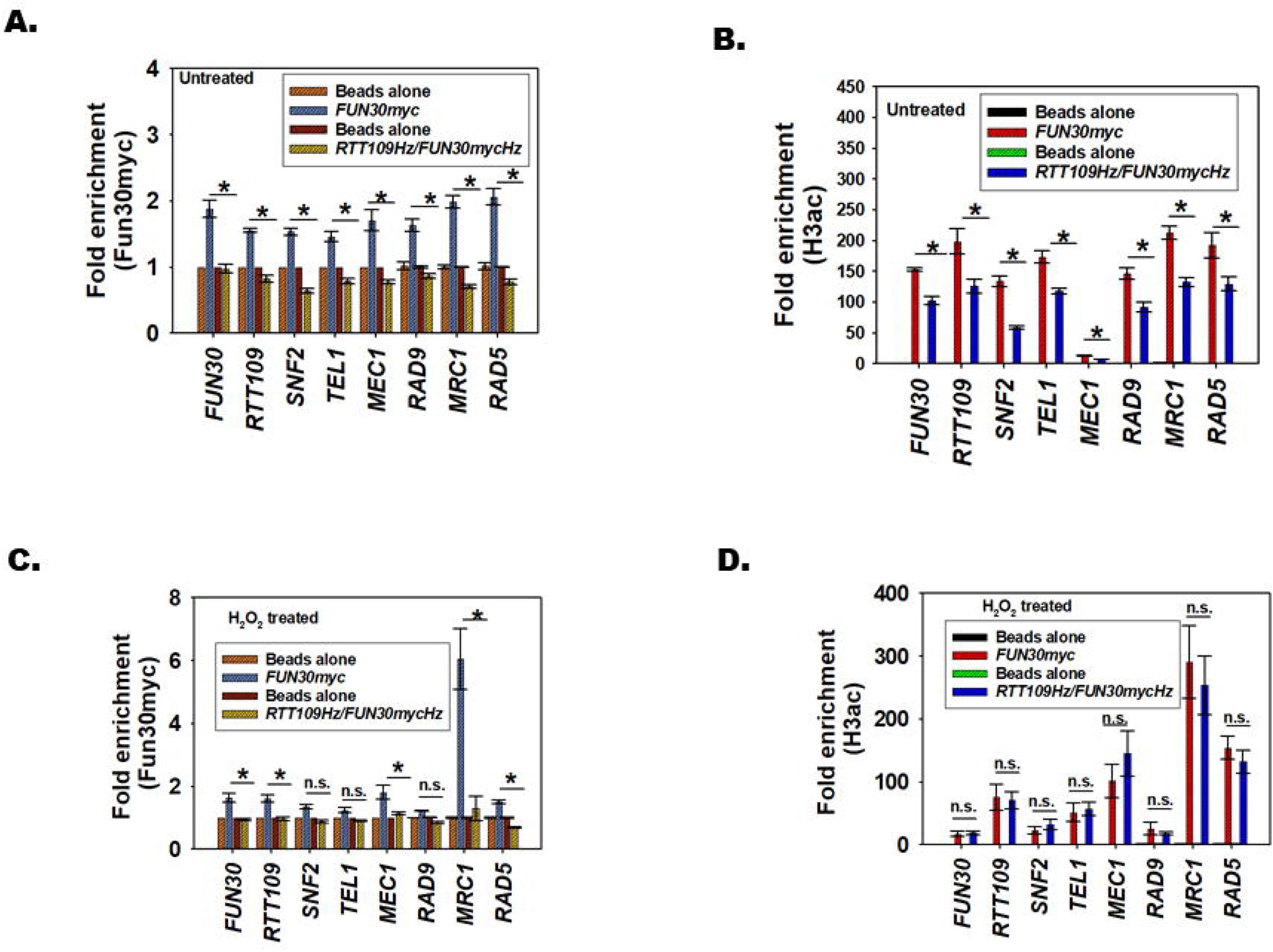
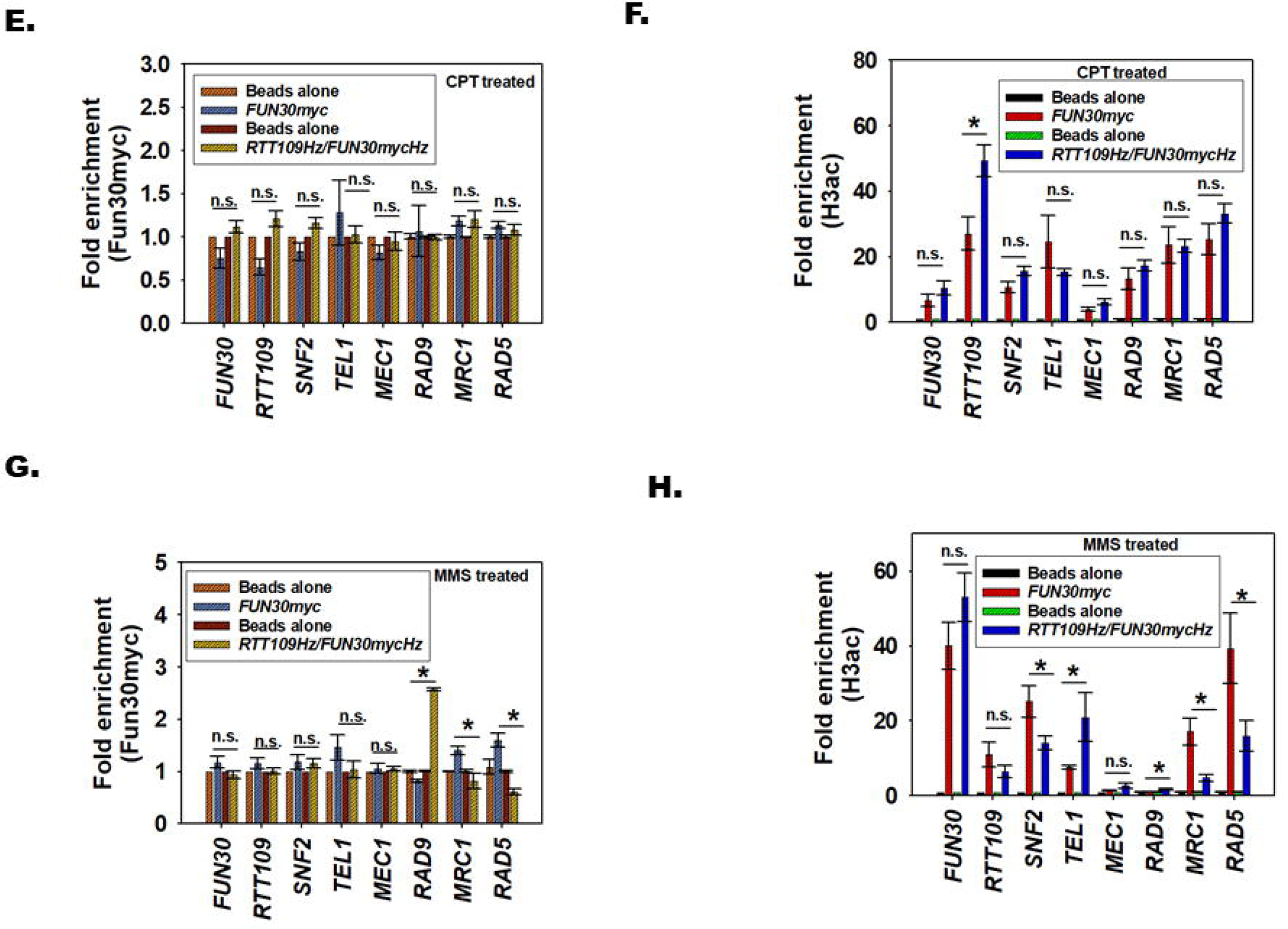
The response to DNA damage is regulated by the occupancy of Fun30 and H3ac on *RAD9*, *MRC1*, and *RAD5* genes. (A). The occupancy of Fun30myc was analysed on *FUN30*, *RTT109*, *SNF2*, *TEL1*, *MEC1*, *RAD9*, *MRC1*, and *RAD5* in *FUN30myc* and *RTT109Hz/FUN30mycHz* in untreated cells. (B). The occupancy of H3ac was analysed on *FUN30*, *RTT109*, *SNF2*, *TEL1*, *MEC1*, *RAD9*, *MRC1*, and *RAD5* in *FUN30myc* and *RTT109Hz/FUN30mycHz* in untreated cells. (C). The occupancy of Fun30myc was analysed on *FUN30*, *RTT109*, *SNF2*, *TEL1*, *MEC1*, *RAD9*, *MRC1*, and *RAD5* in *FUN30myc* and *RTT109Hz/FUN30mycHz* in cells treated with H_2_O_2_ (7.5 mM). (D). The occupancy of H3ac was analysed on *FUN30*, *RTT109*, *SNF2*, *TEL1*, *MEC1*, *RAD9*, *MRC1*, and *RAD5* in *FUN30myc* and *RTT109Hz/FUN30mycHz* in cells treated with H_2_O_2_ (7.5 mM). (E). The occupancy of Fun30myc was analysed on *FUN30*, *RTT109*, *SNF2*, *TEL1*, *MEC1*, *RAD9*, *MRC1*, and *RAD5* in *FUN30myc* and *RTT109Hz/FUN30mycHz* in cells treated with CPT (100 μM). (F). The occupancy of H3ac was analysed on *FUN30*, *RTT109*, *SNF2*, *TEL1*, *MEC1*, *RAD9*, *MRC1*, and *RAD5* in *FUN30myc* and *RTT109Hz/FUN30mycHz* in cells treated with CPT (100 μM). (G). The occupancy of Fun30myc was analysed on *FUN30*, *RTT109*, *SNF2*, *TEL1*, *MEC1*, *RAD9*, *MRC1*, and *RAD5* in *FUN30myc* and *RTT109Hz/FUN30mycHz* in cells treated with MMS (0.02%). (H). The occupancy of H3ac was analysed on *FUN30*, *RTT109*, *SNF2*, *TEL1*, *MEC1*, *RAD9*, *MRC1*, and *RAD5* in *FUN30myc* and *RTT109Hz/FUN30mycHz* in cells treated with MMS (0.02%). The data is presented as average ± s.e.m. of three biological replicates. Star indicates p <0.05.

When cells were treated with H_2_O_2_, the occupancy of Fun30 decreased on *FUN30*, *RTT109*, *MEC1*, *MRC1*, and *RAD5* promoters in the mutant strain as compared to the wild type strain (Fig. 6C). The H3ac occupancy did not alter between the mutant and wild type strain on these promoters (Fig. 6D) indicating that Fun30 is the primary driver for the alteration of gene expression observed during oxidative stress. Thus, *FUN30* but not *RTT109* plays an important role in the response the cell mounts to oxidative stress.

On treatment with CPT, as expected, the occupancy of Fun30 and H3ac was similar in the wild type and mutant strain leading to no change in the gene expression between the two strains in the presence of this genotoxic agent (Fig. 6E and F).

Finally, when cells were treated with MMS, the occupancy of Fun30 and H3ac increased on *RAD9* promoter in the mutant strain as compared to the wild type strain (Fig. 6G and H). Concomitantly, the occupancy of both Fun30 and H3ac decreased on *MRC1* and *RAD5* promoters in the mutant strain as compared to the wild type strain (Fig. 6G and H) leading to increased expression of *RAD9* and decreased expression of *MRC1* and *RAD5*. Thus, on treatment with alkylating agent both Fun30 and H3ac epigenetically modulate the expression of the DNA damage response genes.

#### Single copy deletion of RAD9 in RTT109Hz/FUN30Hz rescues the resistance to MMS

In presence of MMS, *RAD9* is upregulated while *RAD5* and *MRC1* are downregulated in *RTT109Hz/FUN30Hz* as well as *RTT109Hz/FUN30mycHz* as compared to the wild type strains.

In *S. cerevisiae*, deletion of both Rad5 and Fun30 results in resistance to MMS that can be rescued by overexpression of *RAD5* (38). To understand whether this pathway is operational in *C. albicans* also, *RAD5* was ectopically overexpressed in *RTT109Hz/FUN30Hz* mutant to generate *RTT109Hz/FUN30Hz/RAD5OE-URA3* strain. As *URA3* was used as a selection marker for making *RTT109Hz/FUN30Hz/RAD5OE-URA3* strain and overexpression of this gene is known to affect growth, therefore, *URA3* gene was incorporated in both BWP17 and *RTT109Hz/FUN30Hz* strain, creating BWP17-*URA3* and *RTT109Hz*/*FUN30Hz-URA3* strains. In addition, *RAD5* was also overexpressed in BWP17 (*RAD5OE-URA3*) to determine the effect of overexpression of this gene on growth and drug sensitivity in normal cells.

Quantitative PCR indicated that *RAD5* transcript was overexpressed in *RTT109Hz/FUN30Hz/RAD5OE-URA3* strain as well as in *RAD5OE-URA3* (Fig. 7A and Supplementary Fig. 6A). Plate assays showed that *RTT109Hz/FUN30Hz/RAD5OE*-*URA3* mutant was slow growing as compared to BWP17-*URA3* as well as *RTT109Hz/FUN30Hz-URA3* strains in YEPD plate in the absence of MMS (Fig. 7B). However, *RAD5OE*-*URA3* did not show any growth difference as compared to BWP17-*URA 3* (Supplementary Fig. 6B). In the presence of MMS, the growth of the *RTT109Hz/FUN30Hz/RAD5OE-URA3* was slow as compared to *RTT109Hz/FUN30Hz-URA3* mutant but it was impossible to determine whether this slow growth was because of rescue of the phenotype or was it because of the inherent slow growth of the mutant strain (Fig. 7B). It needs to be noted that *RAD5OE*-*URA3* showed greater sensitivity to MMS as compared to BWP17-*URA3* (Supplementary Fig. 6B). Therefore, to understand the effect of *RAD5OE* in *RTT109Hz/FUN30Hz* background, duplication time was calculated for the wild type and the mutant strain in the absence and presence of MMS. In the absence of MMS, *RTT109Hz/FUN30Hz/RAD5OE-URA3* showed 4-fold slower growth as compared to the wild type strain (Fig. 7C and D). However, in the presence of MMS, the growth of *RTT109Hz/FUN30Hz/RAD5OE-URA3* was slower as compared to *RTT109Hz/FUN30Hz-URA3* mutant but faster than BWP17*-URA3* suggesting *RAD5* overexpression can partially rescue the resistant phenotype of *RTT109Hz/FUN30Hz* to MMS (Fig. 7C and D). However, comparison of the growth curve shows that the *RTT109Hz/FUN30Hz/RAD5OE-URA3*was not like that of the BWP17-*URA3* (Fig. 7D).

**Figure 7.**
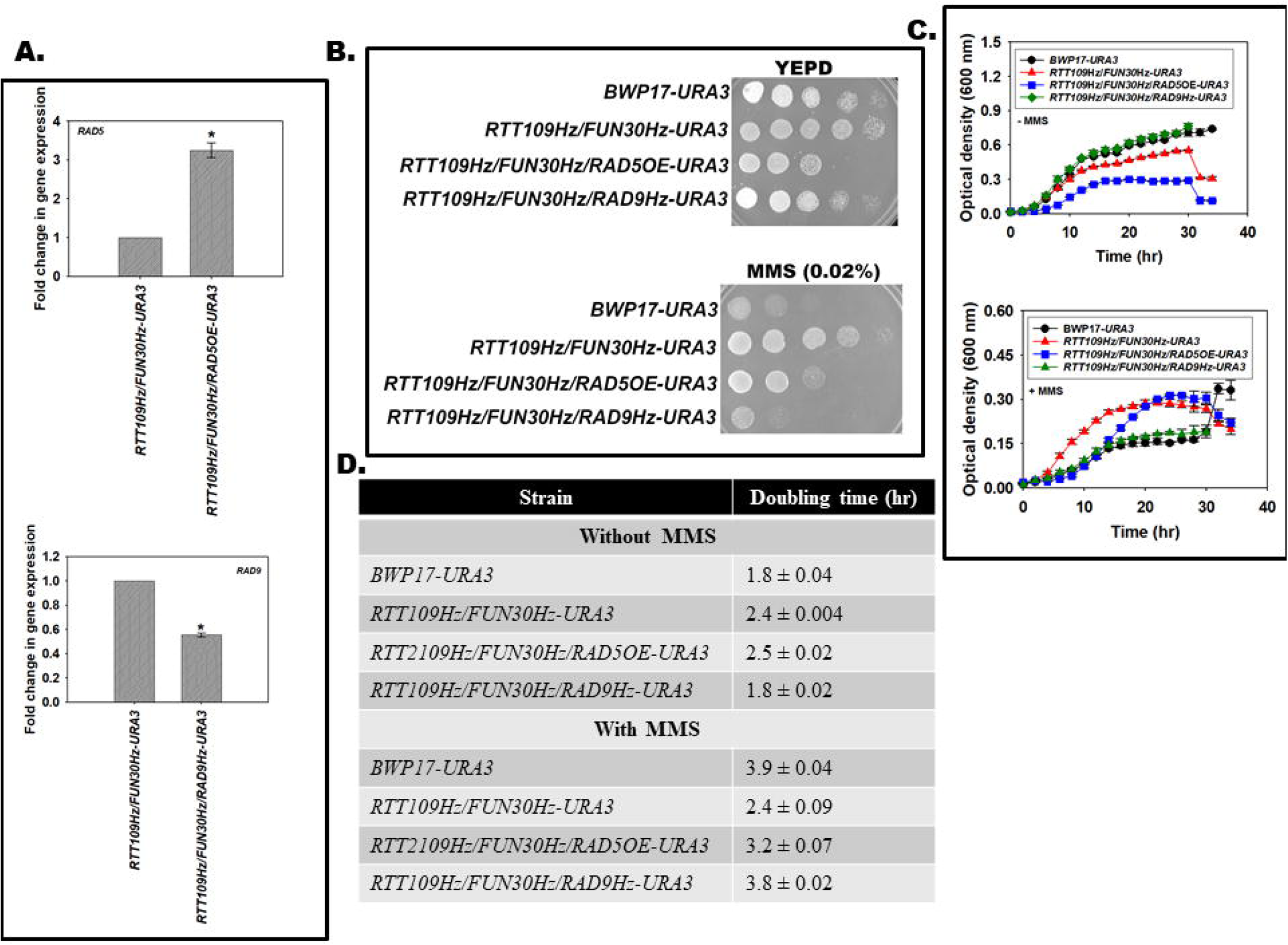
*RAD9* deletion in *RTT109Hz/FUN30Hz* rescues the resistance to MMS. (A). Expression of *RAD5* and *RAD9* was analysed in *RTT109Hz/FUN30Hz-URA3, RTT109Hz/FUN30Hz/RAD5OE-URA3,* and *RTT109Hz/FUN30Hz/RAD9Hz-URA3* strains using qPCR. (B). Sensitivity of BWP17*-URA3*, *RTT109Hz/FUN30Hz-URA3, RTT109Hz/FUN30Hz/RAD5OE-URA3,* and *RTT109Hz/FUN30Hz/RAD9Hz-URA3* to MMS (0.02%) was evaluated using plate assays. (C). Growth curve of BWP17*-URA3*, *RTT109Hz/FUN30Hz-URA3, RTT109Hz/FUN30Hz/RAD5OE-URA3,* and *RTT109Hz/FUN30Hz/RAD9Hz-URA3* in the absence and presence of MMS (0.02%). (D). Table showing the duplication times of the wild type and mutant strains in the absence and presence of MMS.

Next, the role of *RAD9* was investigated. One copy of *RAD9* was deleted creating *RTT109Hz/FUN30Hz/RAD9Hz-URA3*. A control strain *RAD9Hz-URA3* was also created in BWP17 background be deleting one copy of the gene. Quantitative PCR confirmed the expression of the gene was downregulated in both *RTT109Hz/FUN30Hz/RAD9Hz-URA3* and *RAD9Hz-URA3* strains (Fig. 7A and Supplementary Fig. 6C). Plate assays showed that *RTT109Hz/FUN30Hz/RAD9Hz-URA3* and *RAD9Hz-URA3* showed growth similar to BWP17-*URA3* strain on YEPD plate (Fig. 7B and D; Supplementary Fig. 6B). Finally, in the presence of MMS, *RTT109Hz/FUN30Hz/RAD9Hz-URA3* showed sensitivity like the BWP17-*URA3* strain (Fig. 7B). It should be noted that *RAD9Hz-URA3* also shows greater sensitivity as compared to BWP17-*URA3* (Supplementary Fig. 6B). Growth curves confirmed that the duplication time as well as the growth curve of *RTT109Hz/FUN30Hz/RAD9Hz-URA3* and BWP17-*URA3* are similar indicating that *RAD9* overexpression is main reason for the resistance of *RTT109Hz/FUN30Hz* in the presence of MMS (Fig. 7C and D).

## DISCUSSION

DNA damage induced by genotoxic stress activate the DNA damage response pathway. The pathway senses the DNA damage and transduces the message to the DNA damage repair proteins as well as cell checkpoint proteins to initiate DNA damage repair and cell cycle arrest. The Tel1 and Mec1 kinases belonging to the PIKK family are the central transducers of the DNA damage (27). These kinases are activated by autophosphorylation and in turn, phosphorylate the effector proteins. Amongst these are the mediator proteins, Rad9 and Mrc1, that on activation by phosphorylation transduces the signal to Rad53 (36, 39, 40). Rad53, in turn, induces cell cycle arrest (41, 42). In addition to Rad9 and Mrc1, Rad5, an ATP-dependent chromatin remodelling possessing E3 ubiquitination activity, is involved in the replication bypass mechanism when the bases are modified by alkylating agents (37). Studies have also shown that in addition to the role in activation of cell cycle checkpoint activation, Rad9 is required for transcription regulation of DNA damage response proteins (43).

Though extensive studies have shown that the DNA damage response and the DNA damage repair pathways are regulated by post-transcriptional modification, very less information is available about the transcriptional regulation of these pathways.

In this paper, we show that Fun30, an ATP-dependent chromatin remodelling protein, and histone acetylation catalysed by Rtt109, a B-type histone acetyltransferase, transcriptionally co-regulate the response *C. albicans* mounts in the presence of genotoxic stress.

In *C. albicans*, unlike *S. cerevisiae* (8, 44), both copies of *FUN30* could not be deleted. However, in *RTT109* null mutant, Fun30 expression could not be detected by western blot. Therefore, we do not know whether *FUN30* is essential or whether our attempts to delete both copies were unsuccessful.

Like *S. cerevisiae* Fun30, the *C. albicans* protein too possesses DNA-stimulated ATPase activity; however, whether the protein forms homodimer still needs to be determined (10). Using synthetic oligonucleotides, we found that the protein prefers fork DNA as effector unlike ADAAD which prefers stem-loop DNA (45) and Rad54 whose activity is stimulated by double-stranded DNA (46). Binding studies showed that the protein binds to ATP both in the absence and presence of fork DNA; however, the affinity for ATP decreases in the presence of fork DNA. The protein also binds to fork DNA both in the absence and presence of ATP. Further, like in the case of Rad54 (47), the affinity for the DNA does not change in the presence of ATP.

Quantitative PCR and ChIP studies showed that the Fun30 protein in *C. albicans* regulates the expression of *RTT109*, *TEL1*, *MEC1*, *SNF2*, *RAD9*, *MRC1* and *RAD5*. It is also present on its own promoter, possibly regulating itself.

*RTT109* is non-essential and the protein encoded by this gene catalyses the formation of acetylated histones (13, 14). ChIP studies showed that H3ac is present on the promoters of *FUN30, RTT109, TEL1*, *MEC1*, *SNF2*, *RAD9*, *MRC1* and *RAD5*; the occupancy of H3ac decreases on these promoters when both copies of *RTT109* are deleted correlating with decreased expression.

Thus, *FUN30* and *RTT109* both regulate the expression of each other as well as of other DNA damage response genes. ChIP and qPCR data, interestingly, reveal that both Fun30 and H3ac can drive the expression of the DNA damage response genes in the absence of *RTT109* and *FUN30* respectively. However, both Fun30 and H3ac are necessary for mediating the response to genotoxic stress as overexpression of *FUN30* in *RTT109D* strain does not overcome the sensitivity shown by the mutant strain to genotoxic stress.

The double heterozygous mutant *RTT109Hz*/*FUN30Hz* helped to elucidate how these two genes regulate the expression of the DNA damage response genes in the presence of different genotoxic agents. *RTT109Hz*/*FUN30Hz* shows sensitivity to H_2_O_2_, no response to CPT, and resistance to MMS as compared to the wild type cells. qPCR data showed that in the presence of CPT, which induces double-strand break, the expression of the DNA damage response genes are upregulated in *RTT109Hz/FUN30Hz* indicating that the remaining one copy of the two genes is sufficient for the process.

The *RTT109Hz*/*FUN30Hz* mutant is unable to cope with the induction of oxidative stress as the expression of *MEC1*, *RAD9, MRC1* and *RAD5* are downregulated. This downregulation is primarily mediated by the absence of Fun30 as the occupancy of H3ac in the mutant is similar to the wild type. The recruitment of Fun30 to the promoter of the DNA damage response genes in the event of oxidative stress, on the other hand, is impaired leading to downregulation of the genes involved in the DNA damage response pathway resulting in the inactivation of the response/repair. Thus, *RTT109* and *FUN30* are essential for the survival of *C. albicans* in the presence of ROS/oxidative stress.

When cells are treated with MMS, the DNA bases (G and A, primarily) are methylated. In the mutant cells, on treatment with MMS, the occupancy of Fun30 and H3ac on the promoter of *RAD5* decreases leading to reduced expression of Rad5 protein. On the other hand, the occupancy of Fun30 and H3ac increased on *RAD9* promoter correlating with increased expression of this gene.

Methylation of bases results in the stalling of the replication fork and Rad5 plays an important role in the bypass of the stalled replication fork either by recruiting DNA pol η (TLS pathway) or by polyubiquitinating PCNA and bypassing the replication fork (48). By mechanism possibly involving Rad51 and HR mediated repair, deletion of both Fun30 and Rad5 in *S. cerevisiae* has been shown to result in resistance to MMS (38). In *RTT109Hz*/*FUN30Hz* mutant cells, the expression of *RAD5* and *MRC1* is downregulated but not completely abolished. It is possible that reduced expression of *RAD5* results in the bypass TLS pathway leading to faster DNA replication and thus, more growth. However, FACS studies will need to be done to confirm this hypothesis. It also needs to be noted that overexpression of *RAD5* only partially restores sensitivity to MMS indicating that the reduced expression of this gene might not be the primary driver for resistance to MMS in the *RTT109Hz/FUN30Hz* strain.

Concomitantly, the expression of *RAD9* is upregulated in *RTT109Hz/FUN30Hz* cells on treatment with MMS. Deletion of one copy of *RAD9* in *RTT109Hz*/*FUN30Hz* led to restoration of sensitive phenotype in the presence of MMS. Rad9 is known to inhibit DNA end-resection and studies in *S. cerevisiae* has shown that Fun30 inhibits Rad9 to promote DNA end-resection during DNA double-strand break repair (49). Both Fun30 and Rad9 are known to play a role in resistance to MMS; however, the crosstalk between *RTT1109*, *FUN30* and *RAD9* in *C. albicans* in determining the response to MMS needs to be further studied.

Thus, our studies show that the DNA damage response as well as the repair pathway are epigenetically modulated in *C. albicans*.

## MATERIALS AND METHODS

### Chemicals

All chemicals used in this study were of analytical grade and were purchased either from Thermo Fisher Scientific (USA), or Merck (India), or HiMedia (India), or SRL (India), or Sigma-Aldrich (USA). The media for growing cells were purchased from HiMedia (India). Restriction enzymes and T4 DNA ligase were purchased from Thermo Fisher Scientific (USA), Merck (USA) and New England Biolabs (USA). TA cloning kit was purchased from MBI Fermentas (USA). Gel extraction kit was purchased from Qiagen (Germany). SYBR Green was purchased from Kapa Biosystems (Switzerland). Bradford dye for protein estimation was purchased from Sigma-Aldrich (USA). All the primers were synthesized by GCC Biotech (India) (Table S1).

### Plasmids and strains

*E.coli* strain JM109 (Merck, India) was used for cloning *RTT109, FUN30* and *RAD5*. Two *C. albicans* strains BWP17 and SN152 were used in this study and all mutants were made in these backgrounds. BWP17 was a kind gift from Prof. Aaron P. Mitchell, Department of Biological Sciences, Carnegie Mellon University, Pittsburgh, PA, USA. SN152 strain was a kind gift from Prof. Natarajan, JNU and BWP17-*URA3* strain was a kind gift from Prof. S.S.Komath, JNU. The plasmids and strains used in this study are listed in Table S2 and Table S3 respectively.

*C*. *albicans* strains were cultured in Yeast Extract-Peptone-Dextrose (YEPD) media or in synthetic dextrose (SD) minimal media. Ura^-^ strains were cultured in SD media supplemented with 60 μg/ ml uridine. His^-^ strains or Arg^-^ strains were cultured in SD medium supplemented with 85.6 μg/ml histidine or arginine. Transformations were performed using Lithium acetate method (50).

### Antibodies

Antibodies of anti-H3 (Cat# 96C10) was purchased from Cell Signalling Technologies (USA), anti-H3K56ac (Cat# 76307) was purchased from Abcam (UK), anti-c-Myc (Cat# ITG0001) was purchased from Immunotag (USA) and anti-G6PDH (Cat# A9521) was purchased from Sigma-Aldrich (USA).

### Generation of heterozygous and deletion mutants

PCR-mediated gene disruption strategy was used for making all the mutants in *C. albicans*. Heterozygous mutants of *RTT109*, *FUN30* and *VPS75* of were made using the selection marker *HIS1*. Heterozygous mutant of *RAD9* was made using the selection marker *URA3*. For making *RTT109* deletion mutant, the second copy of *RTT109* was replaced with *ARG4* selection marker and transformed in *RTT109Hz* background. For *RTT109Hz/FUN30Hz* mutant, one copy of *RTT109* was replaced with *HIS1* followed by replacing one copy of *FUN30* with *ARG4* selection marker. *RTT109Hz/FUN30/Hz/RAD9Hz-URA3* was created by replacing *RAD9* with *URA3* selection marker in *RTT109Hz/FUN30Hz* background. The *HIS1, ARG4* and *URA3* were amplified by PCR using primers that included flanking gene specific sequence and transformed in the respective *C. albicans* strain. Selection of transformants was done in SD media plate without Histidine or Arginine or Uridine and confirmed by PCR using gene specific flanking primers.

### Generation of overexpression strains

One allele of *RTT109* and *FUN30* were introduced into the *FUN30Hz* and *RTT109D* backgrounds respectively. One allele of *RAD5* was introduced into *RTT109Hz/FUN30Hz* as well as BWP17 strain. For creating the overexpression strains, *RTT109*, *FUN30* and *RAD5* were cloned under *ACT1* promoter in p*ACT1-GFP* vector. *RTT109* was cloned between the HindIII and NheI sites while *FUN30* was cloned between PstI and Nhe1 sites. *RAD5* was cloned into p*ACT1-GFP* vector between the HindIII and NheI sites. The constructs generated were linearized using StuI restriction enzyme and introduced into the respective backgrounds. The transformants were selected on media plate without uridine. The successful transformants were screened by PCR, using gene specific forward primer (FP; Table S1) and locus specific *RPS1* reverse primer (RP; Table S1) to confirm the integration at *RPS1* locus. *URA3* expressing control strains were made by digesting p*ACT1-GFP* vector, which has *URA3* promoter, with Stu1 and transformed.

### Generation of revertant strains

Revertant strains of *RTT109* and *FUN30* in *RTT109* deleted and *FUN30* heterozygous background respectively were performed similarly, as described above.

### Epitope-tagging of endogenous *FUN30*

pADH34 vector, a kind gift of Prof. Johnson, UCSF, was used to add 13X-Myc tag to C-terminus of endogenous *FUN30* of *C. albicans*. Briefly, fusion PCR method was used to generate appropriate double-stranded DNA which was transformed into SN152 strain. The DNA was integrated into the site by homologous recombination and transformed colonies were selected using nourseothricin selection media plate. This strain was termed as *FUN30myc* and all the mutants were made in this background. This strain was used as the control wild type for ChIP, qPCR and plate assays.

### Quantitative PCR (qPCR)

Cells was taken from late log phase secondary culture to isolate the RNA. The cells were collected by centrifugation and then washed twice with DEPC-treated water. The cells were lysed by vortexing with glass beads. TRIzol reagent (Qiagen) was added to the cell pellet to extract the total RNA. cDNA was prepared using 2 µg of total RNA. The transcript levels of different genes were quantified by amplifying with qPCR primers (Table S4) using SYBR Green PCR master mix. *GAPDH* transcripts levels were used as internal control for all the experiments.

### Chromatin Immunoprecipitation (ChIP) assay

Briefly, cells from late log phase secondary culture was harvested after fixing with formaldehyde (1% final concentration) for 20 min. Then glycine (25 mM) was added to quench the formaldehyde and the cells were incubated for 10 min. Following this step, the cells were treated with 5 U lyticase enzyme for 2 hr. Subsequently, the cells were pelleted down, washed and lysed in lysis buffer (50 mM HEPES, pH 7.5; 140 mM NaCl; 1 mM EDTA; 1% Triton X-100; 1 mM PMSF) with glass beads and vortexed. The supernatant was sonicated using a water bath sonicator (five cycles of 15 sec ON and 15 sec OFF each). After sonication, chromatin of different mutants was incubated overnight with respective antibodies. Then, protein G agarose beads were added for binding with the antibody. The beads were subsequently washed with lysis buffer, high-salt buffer (lysis buffer with 500 mM NaCl), wash buffer (10 mM Tris-Cl, pH 8.0; 250 mM LiCl; 1 mM EDTA; 0.5% NP-40) and Tris-EDTA buffer. Finally, elution buffer (1.0% SDS; 100 mM NaHCO_3_) was added to elute the protein-DNA complex and the protein was digested using proteinase K. The DNA was extracted using phenol: chloroform mixture (1:1) and then precipitated with iso-propanol. The ChIP samples were analyzed by qPCR using promoter region specific primers (Table S5).

### Western blots

Cells were taken from 8 hr secondary culture at 30°C and lysates were prepared in ice-cold lysis buffer (50 mM Tris-Cl, pH 7.5; 150 mM NaCl; 50 mM NaF; 1% Triton X-100; 0.1% SDS; 3 mM PMSF) using glass beads. The lysate was centrifuged at 16000 RCF and supernatant were taken to measure the protein concentration. The protein samples were separated by SDS-PAGE. Protein was then transferred from gel to a PVDF membrane. Membrane blocking was performed with 5% skimmed milk in PBS for 1hr. The blot was then probed with respective primary antibody at 4 °C for overnight followed by washing three times with PBST (PBS + 0.05% Tween 20). The blots were incubated with appropriate secondary antibodies. Subsequently, the blots were washed thrice with PBST (10 min per wash) and developed using chemiluminescence.

### Immunofluorescence

Cells from secondary culture were fixed with 37% formaldehyde. After washing the cell pellet with 1X PBS, 5U lyticase treatment was given to digest the cell wall. Triton X-100 was used to permeabilize the cells. Cells were resuspended in blocking solution (1% BSA in 1X PBS). Primary c-Myc antibody was added in a dilution of 1: 100 to the cells and then incubated overnight at 4 °C. The cells were pelleted, washed with PBS, and resuspended in appropriate secondary antibody solution. Hoechst dye was added to it and incubation was continued for 30 min. The cells were pelleted, washed with PBS and mounted on glass slides. The slides were analysed Nikon A1R confocal microscope.

### Growth rate analysis

A primary culture was grown in 10 ml YEPD media at 30°C. 0.1 OD cells from the primary culture was added in secondary culture media and incubated at 30°C, 220 rpm. OD at A_600nm_ was monitored with the aliquots taken from secondary culture every 2 hr until saturation was reached. The growth curve was obtained from the OD values and doubling time was calculated.

### Plate assays

Cells from secondary culture of different mutants were measured at 0.1 OD at A_600nm_ and diluted accordingly with 0.9% saline. A total of five samples of fivefold serial dilution were prepared and spotted on control plates as well as different treated plates. Images were taken for interpretation of results. All plate assays were repeated at least 3 times and the best representative image is shown in the manuscript.

### Overexpression and purification of Fun30

*FUN30* from *C. albicans* was cloned in pET-21c (+) vector between NheI and XhoI. Plasmid DNA was isolated from the transformant and transformed into Rosetta (DE3) cells (Merck Millipore, USA). For purification, a single colony of the transformed cell was inoculated into 10 ml of LB containing 100 μg/ml ampicillin and 50 mg/ml chloramphenicol. The cells were grown for 16 hr at 37°C and 1% (v/v) of the primary inoculum was transferred to fresh LB medium containing ampicillin and chloramphenicol. The cells were grown to an optical density of 0.6, induced with 0.5 mM IPTG and grown at 16°C for 16 hr. The cells were harvested by centrifugation at 5000 rpm for 10 min at 4°C and resuspended in lysis buffer containing 25 mM Tris-Cl pH 7.5, 350 mM NaCl, 5 mM β-mercaptoethanol, 0.5 mM PMSF, and 0.1 mg/ml lysozyme. The cells were homogenized and incubated for 1 hr at 4°C. The cells were lysed by sonication for 5 cycles (15 seconds ON and 45 seconds OFF). After the sonication, cell debris was removed by centrifugation at 12000 rpm for 45 min at 4 °C, and the supernatant was loaded onto Ni^2+^-NTA (GE Healthcare, USA) column pre-equilibrated in equilibration buffer containing 25 mM Tris-Cl pH 7.5, 350 mM NaCl, 5 mM β-mercaptoethanol, and 0.5 mM PMSF. The column was washed, and the bound protein was eluted with buffer containing 200 mM imidazole, 25 mM Tris-Cl pH 7.5, 350 mM NaCl, 5 mM β-mercaptoethanol, and 0.5 mM PMSF. The fractions containing the protein were pooled and imidazole was removed by dialyzing the protein against buffer containing 25 mM Tris-Cl pH 7.5, 100 mM NaCl, 15% (v/v) glycerol, and 5 mM β-mercaptoethanol. Further purification, if necessary, was done using DE52 column. The concentration of the purified protein was determined using Bradford reagent.

### ATPase activity

The ATPase activity of the Fun30 was estimated using NADH coupled oxidation assay. For this assay, 0.2 μM of protein was added in 1X REG buffer containing 25 mM Tris-OAc pH 7.5, 6 mM MgOAc, 60 mM KOAc, 5 mM β-mercaptoethanol, 1.4 mg/ml phosphoenol pyruvate (PEP) and 10 units each of pyruvate kinase (PK) and lactate dehydrogenase (LDH) in a 250 μl reaction volume. ATP and NADH were added to final concentrations of 2 mM and 0.1 mg/ml, respectively. The reaction mixture was incubated for 60 min at 30°C. The change in absorbance was recorded at 340 nm and the concentration of NADH oxidized was calculated using molar extinction coefficient of NADH as 6.3 mM^−1^. For DNA-stimulated ATPase activity, 500 nM concentration of single-stranded DNA (ssDNA), double-stranded DNA (dsDNA), stem-loop DNA (slDNA), fork DNA, and replication fork DNA were used. The sequence of these DNA molecules is provided in Table S6.

### Fluorescence studies

Ligand binding study based on tryptophan specific excitation were conducted using fluorescence spectrophotometer (Cary Eclipse fluorimeter). The concentration of Fun30 used for fluorescence studies was 0.5 μM. All the experiments were done in room temperature in the buffer containing 50 mM Tris-SO_4_ pH 7.5, 5 mM β-mercaptoethanol and 1 mM MgSO_4_. The excitation wavelength was 295 nm, and the emission wavelength was 340 nm. The slit widths were 5 and 10 nm, respectively. The spectra obtained were corrected for dilution and the inner filter effect was negligible. The binding data obtained were fit to a one-site saturation for the interaction of the ligand with the protein. The *K*_d_ was calculated using the equation

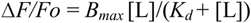

Where *B*_max_ is the maximal binding, [L] is the ligand concentration, and *K*_d_ is the dissociation constant.

### Phylogenetic, Domain, and motif analysis

Phylogenetic analysis of uncharacterised Fun30 protein sequence from *C. albicans* was done using www.phylogeny.fr/version2_cgi/ using default settings. The protein sequences of different ATP-dependent chromatin remodelling proteins from various organisms belonging to different families were extracted from NCBI. Multiple sequence alignment was performed using T-COFFEE Multiple sequence alignment (http://tcoffee.crg.cat/). The domain architecture of Fun30 and ΔNFun30 from *C. albicans* was created using https://prosite.expasy.org/mydomains/ website in its default settings. To identify the conserved motif sequences in Fun30 from *C. albicans*, motif analysis was done using http://tcoffee.crg.cat/apps/tcoffee/do:psicoffee website.

For phylogenetic and motif analysis, orf19.6291 protein from *C. albicans*, Rad5 from *S. cerevisiae*, Rad16 from *S. cerevisiae*, Lodestar from *D. melanogaster*, Etl1 from *M. musculus* and SMARCAD1 from *H. sapiens*, Fun30 from *S. cerevisiae*, Fft1, Fft2, and Fft3 from *S. pombe*, Iswi from *D. melanogaster*, Snf2 from *S. cerevisiae*, BRG1, BRM, and CHD7 from *H. sapiens*, Mi-2 from *D. melanogaster*, CHD1 from *S. cerevisiae*, EP400 from *H. sapiens* and *M. musculus*, SWR1 from *S. cerevisiae* and *C. elegans*, Ino80 from *H. sapiens*, *S. cerevisiae* and *C. elegans*, ERCC6 from *H. sapiens*, ATRX from *H. sapiens*, Rad54 from *S. cerevisiae*, SMARCAL1 from *H. sapiens* and Mot1 from *S. cerevisiae* were used.

## AUTHOR CONTRIBUTIONS

Conceptualization, R.M., P.G., P.K.M., and M.G.; Methodology, R.M., P.G., P.K.M., and M.G.; Investigation, P.G., P.K.M., M.G., D.T.H., H.B., A.G., and S.D.; Resources - R.M.; Writing - original draft, R.M., P.K.M., and P.G.; Writing-review and editing, R.M., P.G., and P.K.M.,; Funding acquisition, R.M.; Supervision, R.M.

## ACKNOWLEDGEMENTS

The authors would like to thank the Central Instrumentation facility, School of Life Sciences for confocal microscope and Fluorescence Spectrophotometer.

