## Supplementary Tables and figure legends for "RTT109 and Fun30 proteins mediate epigenetic regulation of the DNA damage response pathway in *C. albicans*"

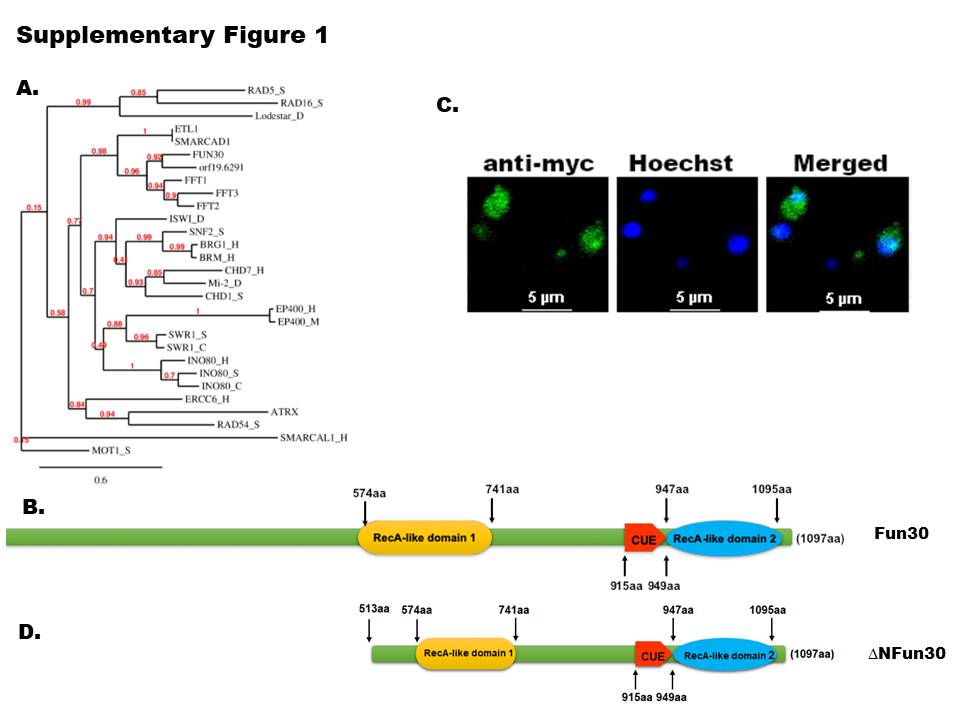

**Supplementary Figure 1. The** **orf19.6291 encodes for Fun30, a member of the ATP-dependent chromatin remodelling protein family. A.** Phylogenetic analysis of orf19.6291 protein from *C. albicans* was done with Rad5 from *S. cerevisiae* (RAD5_S), Rad16 from *S. cerevisiae* (RAD16_S), Lodestar from *D. melanogaster* (Lodestar_D), Etl1 from *M. musculus* and SMARCAD1 from *H. sapiens*, Fun30 from *S. cerevisiae*, Fft1, Fft2, and Fft3 from *S. pombe* (FFT1, FFT2, and FFT3 respectively), Iswi from *D. melanogaster* (ISWI_D), Snf2 from *S. cerevisiae* (SNF2_S), BRG1, BRM, and CHD7 from *H. sapiens* (BRG1_H, BRM_H, CHD7_H respectively), Mi-2 from *D. melanogaster* (Mi-2_D), CHD1 from *S. cerevisiae* (CHD1_S), EP400 from *H. sapiens* and *M. musculus* (EP400_H and EP400_M respectively), SWR1 from *S. cerevisiae* and *C. elegans* (SWR1_S and SWR1_C respectively), Ino80 from *H. sapiens*, *S. cerevisiae* and *C. elegans* (INO80_H, INO80_S, and INO80_C respectively), ERCC6 from *H. sapiens* (ERCC6_H), ATRX from *H. sapiens*, Rad54 from *S. cerevisiae* (RAD54_S), SMARCAL1 from *H. sapiens* (SMARCAL1_H) and Mot1 from *S. cerevisiae* (MOT1_S). **B.** Domain architecture of Fun30 (orf19.6291) from *C. albicans*. **C.** Localization of Fun30 protein from *C. albicans* was monitored using immunofluorescence. Fun30 was tagged with myc and localization was monitored using anti-myc antibody. The nucleus was stained using Hoechst. **D.** Domain architecture of ΔNFun30.

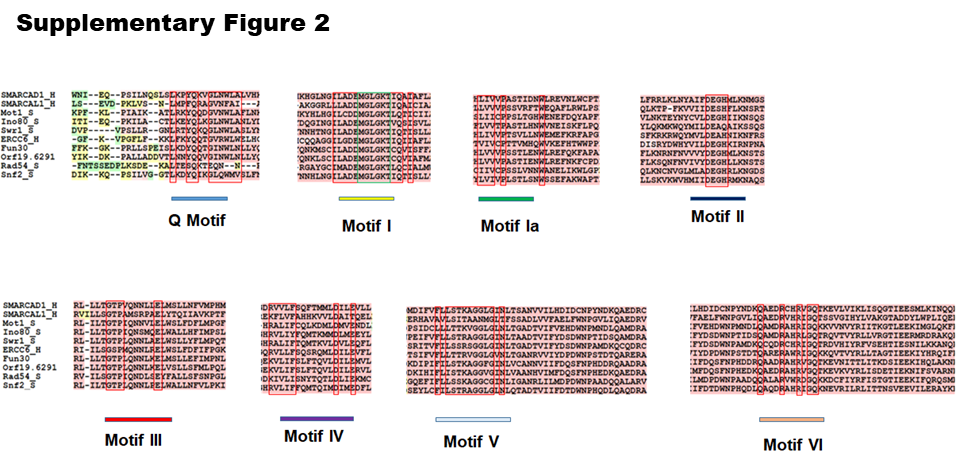

**Supplementary Figure 2. Fun30 from *C. albicans* possesses the seven helicase motifs**. Multiple sequence alignment of SMARCAD1 from *H. sapiens* (SMARCAD1_H), SMARCAL1 from *H. sapiens* (SMARCAL1_H), Mot1 (Mot1_s), Ino80 (Ino80_s), Swr1 (Swr1_s) from *S. cerevisiae*, ERCC6 from *H. sapiens* (ERCC6_H), Fun30 from *S. cerevisiae* (Fun30), orf19.6291 from *C. albicans* (orf19.6291), Rad54 (Rad54_s) and Snf2 (Snf2_S) from *S. cerevisiae* was performed to identify the seven helicase motifs. The presence of these seven motifs in orf19.6291 further establishes that the protein encoded by this ORF is a member of the ATP-dependent chromatin remodelling protein family.

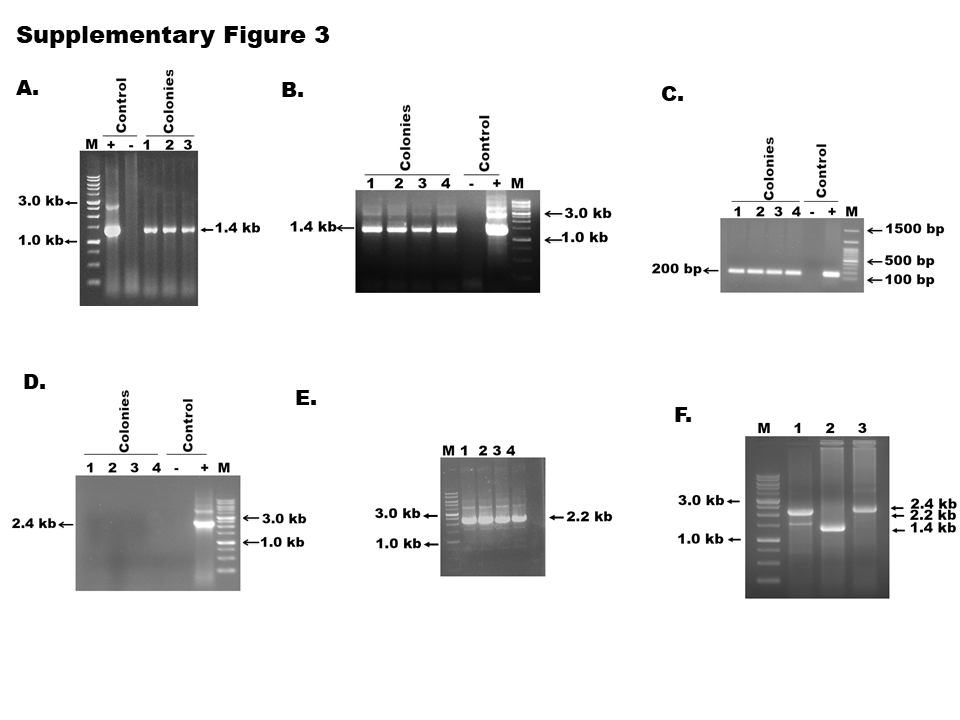

**Supplementary Figure 3. The *FUN30Hz* mutant was confirmed using PCR. A.** One copy of *FUN30* was deleted in *FUN30myc strain*. Three colonies (# 1, 2, and 3) were screened using*HIS1*cassette specific primers and all the colonies gave the expected 1.4 kb product. A positive control containing only the purified pmCherry-*HIS1* plasmid and a negative control lacking both the control plasmid and genomic DNA were included in the experiment. **B.** One copy of *FUN30* was deleted in BWP17 strain. Four transformants (#1, 2, 3, 4) were screened using *HIS1*cassette specific primers and the appearance of 1.4 kb band indicated that one copy of the gene had been successfully deleted. **C.** The transformants were further confirmed using *FUN30* specific product. The appearance of a 200 bp product confirmed *FUN30Hz.* **D.** Attempts were made to delete the second copy of *FUN30* in *FUN30Hz* mutant made in BWP17 background. Four colonies (#1,2, 3,4) were screened using *ARG4*cassette specific primers. The expected 2.4 kb band (positive control using purified pRS-*ARG4* plasmid) was not observed in the four colonies indicating that the second copy of the gene had not been deleted. **E.** Overexpression of *FUN30* in *FUN30Hz* strain was done by transforming the strain with p*ACT1-FUN30* construct. The appearance of a 2.2 kb band after amplification with *FUN30* specific forward primer and *RPS1* specific reverse primer indicating positive transformants. The strain was named as *FUN30Hz*/*FUN30OE-URA3*. **F.** The second copy of *FUN30* was deleted in *FUN30Hz*/*FUN30OE-URA3.* Lane M – Marker. Lane 1- The overexpressed copy of *FUN30* was amplified using *FUN30* specific forward primer and *RPS1* specific reverse primer. PCR amplification gave the expected 2.2 kb band. Lane 2- Deletion of one copy of *FUN30* was confirmed by the appearance of 1.4 kb band amplified using primers specific for *HIS1* marker. Lane 3-Deletion of the second copy of *FUN30* was confirmed by the appearance of 2.4 kb band amplified using primers specific for *ARG4* marker.

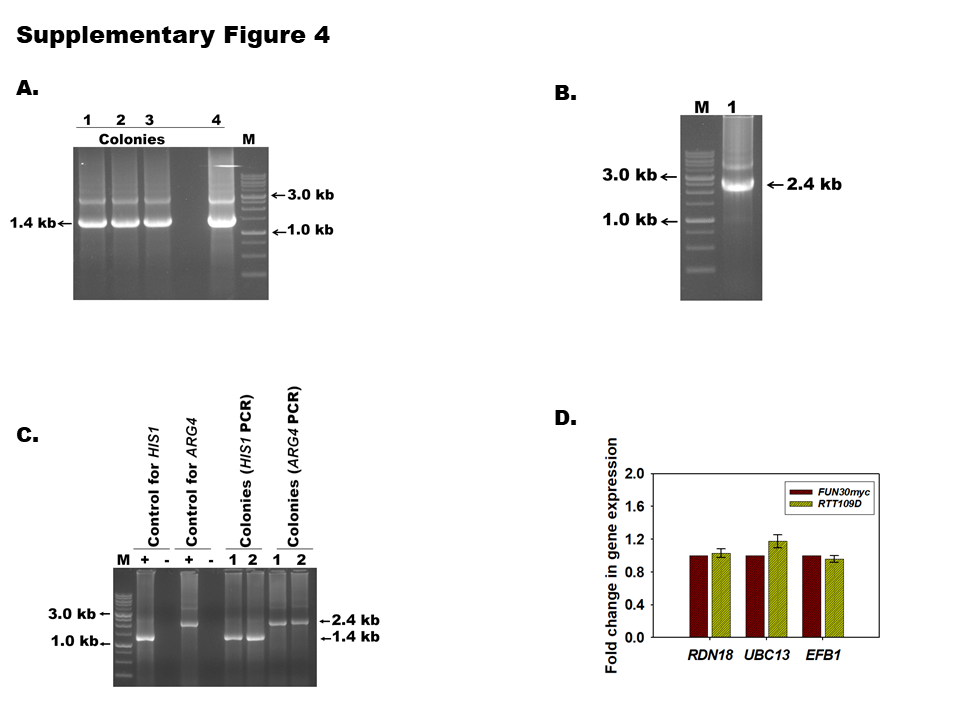

**Supplementary Figure 4. The *RTT109Hz* and *RTT109D* mutants was confirmed using PCR. A.** One copy of *RTT109* was deleted in BWP17 strain. Four colonies (# 1, 2, 3 and 4) were screened using *HIS1* specific primers and all the colonies gave the expected 1.4 kb product. **B.** The deletion of the second copy of *RTT109* in *RTT109Hz* strain was confirmed by PCR using *ARG4* specific primers. The appearance of the expected 2.4 kb band confirmed the mutant strain. **C.** Deletion of the second copy of *RTT109* in *RTT109Hz* strain made in *FUN30myc* background was confirmed by PCR using *HIS1* and *ARG4* cassette specific primers. Two colonies (1 and 2) were screened. The appearance of the expected 1.4 kb and 2.4 kb confirmed the mutants. **D.** Expression of *RDN18*, *UBC13* and *EFB1* was analysed by qPCR in *FUN30myc* and *RTT109D* strain

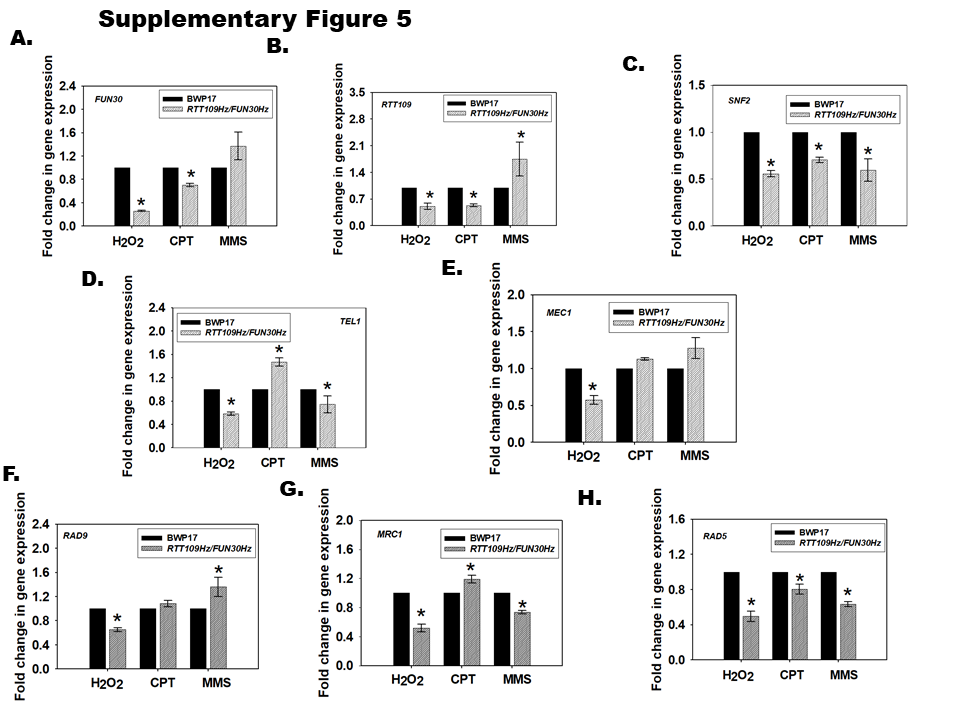

**Supplementary Figure 5. Rtt109 and Fun30 co-regulate the transcription of genes involved in DNA damage response pathways.** Expression of **A.** *FUN30*; **B.** *RTT109;* **C.** *SNF2*; **D.** *TEL1*; **E.** *MEC1*; **F.** *RAD9*; **G.** *MRC1*; **H.** *RAD5* was analysed in BWP17 and *RTT109Hz/FUN30Hz* strains in the presence of H_2_O_2_ (7.5 mM), CPT (100 μM) and MMS (0.02%).

The qPCR data is presented as average ± s.e.m. of three biological replicates. Star indicates p <0.05.

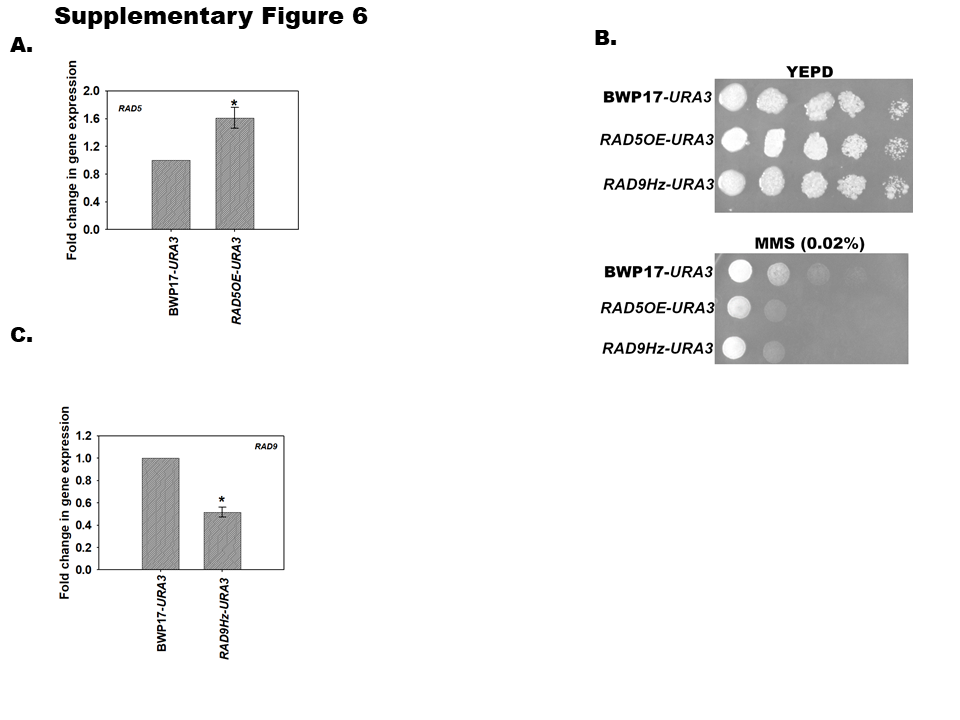

**Supplementary Figure 6. Overexpression of *RAD5* and single copy deletion of *RAD9* leads to sensitivity to MMS.** **A.** Expression of *RAD5* in BWP17*-URA3* and *RAD5OE-URA3* strain was estimated using qPCR. **B.** Plate assays to analyse the sensitivity of *RAD5OE-URA3* and *RAD9Hz-URA3* to MMS. **C.** Expression of *RAD9* in BWP17*-URA3* and *RAD9Hz-URA3* strain was analysed by qPCR. The qPCR data is presented as average ± s.e.m. of three biological replicates. Star indicates p <0.05.

**Supplementary tables**

**Table S1: List of primers used for constructing mutants.**

| **Primer name** | **Primer sequence (5'⭢ 3')** |
| --- | --- |
| RTT109-HindIII-FP | GCGAAGCTTGATTCTTAAAATCGCGTCCTTTCA |
| RTT109-NheI-RP | GCGGCTAGCCTATTTTGATTTTCTATAATTACT |
| RTT109-HIS1-FP | ATGCTTCCTCCAGATATATTACAAAATGGTGAATTTGAAA  CCATTTACTTCCAGACAAATCGGGGATCCTGGAGGATGAG |
| RTT109-HIS1 -RP | CTATTTTGATTTTCTATAATTACTTACAAAATTACTAACT  TCAACACGATCACTAAAATCCGGAATATTTATGAGAAACT |
| RTT109-ARG4-FP | ATGCTTCCTCCAGATATATTACAAAATGGTGAATTTGAAA  CCATTTACTTCCAGACAAATTGTGGAATTGTGAGCGGAAG |
| RTT109-ARG4-RP | CTATTTTGATTTTCTATAATTACTTACAAAATTACTAACTT  CAACACGATCACTAAAATCTTTCCCAGTCACGACGTT |
| FUN30-PstI-FP | GCGCTGCAGATGAGTTGGTTTAGAAGAAATAAAC |
| FUN30-NheI-RP | GCGGCTAGCTCAACTATAAACTATTGACTCTAATG |
| FUN30-HIS1-FP | ATGAGTTGGTTTAGAAGAAATAAACCAACGGAGGAATCA  TCTACAGCTGATCCAAACACACGGGGATCCTGGAGGATGAG |
| FUN30-HIS1-RP | TCAACTATAAACTATTGACTCTAATGTTGAAATCTTTTCG  TCTAACTCTAATTTGTTTCTCGGAATATTTATGAGAGAAACT |
| FUN30-ARG4-FP | ATGAGTTGGTTTAGAAGAAATAAACCAACGGAGGAATCA  TCTACAGCTGATCCAAACACATGTGGAATTGTGAGCGGAAG |
| FUN30-ARG4-RP | TCAACTATAAACTATTGACTCTAATGTTGAAATCTTTTCG  TCTAACTCTAATTTGTTTCTTTTCCCAGTCACGACGTT |
| FUN30-MYC-FP | GAAACAAATTAGAGTTAGACGAAAAGATTTCAACATTAGAGTCAATAGTTTATAGTCGGATCCCCGGGTTAATTAACGG |
| FUN30-MYC-RP | TTACCAAAACAAGCGGGAAAATCAGCTTCAAATGTGTACATTTCTTATTTTGCGGCGGCCGCTCTAGAACTAGTGGATC |
| RAD9-URA3-FP | ATACAATTGCCAGAAACTCAATCTCAAAGCTTATTATATTATGATTCACACTAGAAGGACCACCTTTGATTG |
| RAD9-URA3-RP | TTTGGTGCTATAATTGGGATTATTTCATCTGATTGTTTTTTAGATTCTGATTTGTACAATTCATCCATAC |
| RAD5-HindIII-FP | CCCAAGCTTATGAAAGTTATAAAGAAAAGG |
| RAD5-Nhe1 -RP | CTAGCTAGCCTATTCCTCAAAAAGGATTTG |
| ΔN Fun30-FP | GCTAGCATGGATTTGGACCCCATTGAG |
| ΔN Fun30-RP | CTCGAGACTATAAACTATTGACTC |
| RPS1-RP | AATAGAGAGAAACTATATTATACAC |

**Table S2: List of plasmids used in this study.**

| **Plasmids** | **Reference** |
| --- | --- |
| p*ACT1-GFP* | Alistair Brown, Aberdeen |
| p*ACT1-RTT109* | This study |
| p*ACT1-FUN30* | This study |
| p*ACT1-RAD5* | This study |
| pmCherry*-HIS1* | Victoria et al. (1) |
| pRS*-ARG4* | Wilson et al., 1999 (2) |
| *URA3-*p*MET3-GFP* | Gerami-Nejad et al., 2004 (3) |

**Table S3: List of strains used in this study.**

| **Strains** | **Genotype** | **Referred in manuscript** | **Reference** |
| --- | --- | --- | --- |
| **Mutants made using BWP17 strain** | | | |
| BWP17 | *ura3*::imm434*/ura3*::imm434 *iro1/iro1*::  imm434 *his1*::hisG*/his1*::hisG*arg4/arg4* | BWP17 | Wilson et al., 1999 (2) |
| BWP17*-URA3* | BWP17 with *RPS1*/*rps1Δ::*  *pACT1-GFP::URA3* | BWP17*-URA3* | Jain et al., 2010 (4) |
| *VPS75/vps17*-*URA3* | BWP17with *VPS75/vps75Δ::ARG4* with *RPS1/rps1Δ::pACT1-GFP::URA3* | *VPS75Hz*-*URA3* | This study |
| *RTT109/rtt109* | BWP17 with *RTT109/rtt109Δ::HIS1* | *RTT109Hz* | This study |
| *RTT109/rtt109-URA3* | BWP17-*RTT109* heterozygous with *RPS1/rps1Δ::pACT1-GFP::URA3* | *RTT109Hz-URA3* | This study |
| *rtt109/rtt109*-*URA3* | BWP17-*RTT109* heterozygous *URA3* with *RTT109/rtt109Δ::ARG4* | *RTT109D*-*URA3* | This study |
| *rtt109/rtt109/*  p*ACT1-RTT109* | BWP17-*RTT109* deletion with *RPS1*/*rps1Δ::pACT1-RTT109* | *RTT109D/*  *RTT109OE-URA3* | This study |
| *rtt109/rtt109/*  p*ACT1-FUN30* | BWP17-*RTT109* deletion with *RPS1*/*rps1Δ::pACT1-FUN30* | *RTT109D/*  *FUN30OE-URA3* | This study |
| *FUN30/fun30* | BWP17 with *FUN30/fun30Δ::HIS1* | *FUN30Hz* | This study |
| *FUN30/fun30*-*URA3* | BWP17- *FUN30* heterozygous with *RPS1/rps1Δ::pACT1-GFP::URA3* | *FUN30Hz*-*URA3* | This study |
| *FUN30/fun30/*  p*ACT1-FUN30* | BWP17-*FUN30* heterozygous with *RPS1/rps1Δ::pACT1-FUN30* | *FUN30Hz/*  *FUN30OE-URA3* | This study |
| *FUN30/fun30/*  p*ACT1-RTT109* | BWP17-*FUN30* heterozygous with *RPS1/rps1Δ::pACT1-RTT09* | *FUN30Hz/*  *RTT109OE-URA3* | This study |
| *RTT109/rtt109/*  *FUN30/fun30* | BWP17-*RTT109* heterozygous with *FUN30/fun30Δ::ARG4* | *RTT109Hz/*  *FUN30Hz* | This study |
| *RTT109/rtt109/*  *FUN30/fun30 -URA3* | BWP17-*RTT109* and *FUN30* heterozygous with *RPS1*/*rps1Δ::pACT1-GFP::URA3* | *RTT109Hz/*  *FUN30Hz -URA3* | This study |
| *RTT109/rtt109/*  *FUN30/fun30/*  *RAD9/rad9-URA3* | BWP17-*RTT109* and *FUN30* heterozygous with *RAD9/rad9Δ::URA3* | *RTT109Hz/*  *FUN30Hz/*  *RAD9Hz-URA3* | This study |
| *RTT109/rtt109/*  *FUN30/fun30/*  p*ACT1-RAD5* | BWP17-*RTT109* and *FUN30* heterozygous with *RPS1*/*rps1Δ::pACT1-RAD5* | *RTT109Hz/*  *FUN30Hz/*  *RAD5OE-URA3* | This study |
| **Mutants made using SN152 strain** | | | |
| SN152 | *ura3/*::imm434::*URA3/ura3*::imm434 *iro1::IRO1/iro1*::imm434 *his*1::hisG/*his1*::hisG *leu2/leu2 arg4/arg4* | SN152 | Noble and Johnson, 2005 (5) |
| *FUN30myc* | SN152 with *FUN30*-13X myc-*FLP-SAT1* | *FUN30myc* | This study |
| *RTT109/rtt109* | *FUN30myc* with *RTT109/rtt109Δ::HIS1* | *RTT109Hz* | This study |
| *FUN30myc/fun30* | *FUN30myc* with *FUN30/fun30Δ::HIS1* | *FUN30mycHz* | This study |
| *rtt109/rtt109* | *FUN30myc*-*RTT109* heterozygous with *RTT109/rtt109Δ::ARG4* | *RTT109D* | This study |
| *RTT109/rtt109/*  *FUN30myc/fun30* | *FUN30myc*-*RTT109* heterozygous with *FUN30/fun30Δ::ARG4* | *RTT109Hz/*  *FUN30mycHz* | This study |

**Table S4: List of primers used for qPCR analysis.**

| **Gene** | **Forward primer (5'⭢ 3')** | **Reverse primer (5'⭢ 3')** |
| --- | --- | --- |
| *RTT109* | TCGTTGATTGGATGCTGTAAGG | ACCAGCTTCAACAGGTTCATAA |
| *FUN30* | GTGGAATTGAACCAAGTGTAGCTG | TGAGATTGCCTTCCGTTGTCTC |
| *TEL1* | ATTCTACCAGTTGGCTTGCGA | TCCAAAGTTGTTCTTTCCGGC |
| *MEC1* | GAGACACAGCAAGACCCATTA | CGAGCAACTTGTCATCTTTCAG |
| *SNF2* | TGGAGGAGTATGGTCGTGGT | TCTTCGGCTTGGCTTCCATT |
| *RAD5* | GGCCATACGCATCTCAAACT | CGGCGTTTGGATAATCTTGT |
| *RAD9* | TCAAATTCATTGGTGGCTGA | TTCGTTTTCGTTCTCGTTCA |
| *MRC1* | TGACGAACAAGCCACTCAAG | TCATTTTACGACCACGACGA |
| *UBC13* | TCGGTCCTAATCAATCACCTT | TCTTTCAACACATCCAAACAAA |
| *RDN18* | CCACCACCCACAAAATCAA | CGGCACCTTACGAGAAATCA |
| *EFB1* | GCTTCTGGTTCTGCTGCTG | GCTGGTTTTGGACCTTTAGC |
| *VPS75* | CCAATGTATGCCAAAAGACG | AATGACCTGCCTCATCATCG |
| *GAPDH* | GTCCATCCCACAAGGACTGG | CAACGGAAACATCGGTGGTTGG |

**Table S5: List of primers used for ChIP analysis.**

| **Gene** | **Forward primer (5'⭢ 3')** | **Reverse primer (5'⭢ 3')** |
| --- | --- | --- |
| *FUN30* promoter | AATCAGTTGTATAGGAAGGAGAT | GCAAAAGTTGGTTATTTGTACTAG |
| *SNF2*  promoter | GATGAGGGTGGTTGGAGATAT | GGATGAATACTATGTATTGGGTCG |
| *MEC1*  promoter | ATTATTCAAGAGACCCTTAAATCCC | ATTTCTATCCTATTTCTGAGGAGGG |
| *TEL1*  promoter | GGTAGAGAGAGTGACAGTATCAACT | ATATCTGACGTAGACATGATTACG |
| *RTT109*  promoter | CCATCCCAGTTAATTGTTTACCATCTG | GAATACACCACAATAACAACACTTGCTAAT |
| *RAD5* promoter | CCAACCTTCACTCTAACAAACATTTAG | TGGGAGAAGTGGTTGTTTCTTC |
| *RAD9* promoter | AGTGTGCACAACTTGAATGG | GTGAATTCTTTGATGAGGCAGAT |
| *MRC1*  promoter | GCAGATCCCAATTTGTAACACT | CGTACGGTTTAGCTTAGAAAGAAA |
| *Chr5* intergenic region | ATGGGTGTGCTGCTTTTGTT | TAAGGAAGTTGTCGGGTCGA |
| *GAPDH*  promoter | CAGCTGTTTCAAATCCAGGCT | GTGGTTGAGTGGGTTGGTTG |

**Table S6: Oligonucleotides used for ATPase assays**

| **Oligonucleotide** | **Primer sequence (5'⭢ 3')** |
| --- | --- |
| **ssDNA** | CACGCGAAGTGGACCTCG |
| **dsDNA** | CGCGTTAACGCG  CGCGTTAACGCG |
| **slDNA** | GCGCAATTGCGCTCGACGATTTTTTAGCGCAATTGCGC |
| **Fork DNA** | GCGCAATTGCGCTCGACGATTTTTTTTTTGG  TTTTTTTTTTTTTTTTTTAGCGCAATTGCGC |
| **Replication fork DNA** | CCAAAAAAAAAATC  GCGCAATTGCGCTCGACGATTTTTTTTTTGG  TTTTTTTTTTTTTTTTTTAGCGCAATTGCGC |
